# Verifiable Agentic Science on Public Infrastructure

**DOI:** 10.64898/2026.10.26.761864

**Authors:** Dannon Baker, Scott Cain, Marius van den Beek, John Chilton, Danielle Callan, Steven Weaver, Nate Coraor, Kelsey Beavers, Jeremy Goecks, Michael C. Schatz, Björn Grüning, Sergei L. Kosakovsky Pond, Anton Nekrutenko

**Author notes:** galaxyproject.org (Dannon Baker, Scott Cain, Marius van den Beek, John Chilton, Nate Coraor, Jeremy Goecks, Michael C. Schatz, Björn Grüning, Anton Nekrutenko). usegalaxy.eu (Björn Grüning).

## Abstract

Autonomous artificial intelligence agents are transforming the practice of science. Across every discipline, researchers are handing data analysis to coding agents that can design and execute complex studies from a single prompt. Yet this revolution carries a threat. When agents improvise in local environments, they leave no verifiable record, ultimately risking a wave of irreproducible science. Solving this crisis requires moving agentic science onto public infrastructure where every action leaves an examinable trail. Here we connect autonomous coding agents to shared computing resources, turning ephemeral prompts into auditable scientific records. After showing that 30 unguided local agents splinter into 18 incompatible pipelines on the same dataset, we conduct an agentic reproduction and discovery study on a large scale—reproducing a large, diverse array of studies with every workflow, history, and intermediate dataset publicly accessible. This complete provenance trail exposes unstated modeling choices, reference mismatches, and subsampled tutorial inputs in published studies, and establishes a new foundation where human scientists can understand, steer, and trust AI-driven discovery.

## Introduction

Across all scientific domains, language-model agents now formulate hypotheses, build pipelines, write code, and draft manuscripts [1–6]. Although several groups have built specialized research agents [7], domain-specific wrappers cannot keep pace with general-purpose command-line coding agents. At our own institutions, students, postdocs, and principal investigators—including long-time AI skeptics—now use command-line coding agents as their primary working environment.

Yet running autonomous agents on personal machines is equally thrilling and terrifying. Researchers can launch complex analyses beyond their training in minutes, but three flaws threaten reliability. Unsupervised local agents leave no verifiable provenance, generating disposable scripts and tables that vanish when the session closes. Without curated domain guidance, agents improvise fragile pipelines from unvetted snippets, and local execution hits a hardware and security wall whenever an analysis requires high-memory alignment, large cohorts, or GPU inference on shared clusters. Without workflow, tool, and dataset provenance on shared public infrastructure, autonomous analyses are scientifically unverifiable.

For over twenty years, the Galaxy platform has provided open, reproducible infrastructure to life scientists worldwide [8], offering a natural public plane for agentic science. Agents have been connected to workflow platforms before—such as ClawBio’s Galaxy Bridge over BioBlend and comparable tooling for Nextflow and AWS HealthOmics—yet what has been missing is a harness that keeps a checkable record of that work on free public compute. Any coding agent—or Orbit, our desktop workbench—can connect to Galaxy to draw on thousands of vetted containers, peer-reviewed workflows, and community training tutorials [9–14] on national supercomputers without requiring cluster credentials. Every analytical step, tool version, parameter setting, and intermediate dataset is recorded in a durable, shareable history and visual workflow linked to a research notebook, allowing human scientists to understand, steer, and verify what the agent is doing.

Here we show why autonomous agents need shared public infrastructure and what they can achieve once connected to it. After illustrating how unassisted local agents splinter into conflicting pipelines and disposable scripts on a single dataset, we conduct a large-scale validation and reanalysis of published studies spanning bacterial, fungal, vector, mammalian, parasitic, and viral systems on UseGalaxy.org and national supercomputers. Across spatial transcriptomics, single-cell atlases, bulk RNA-seq, chromatin accessibility, protein structure phylogenomics, and epidemic dating, every reanalysis checks out and leaves a complete, publicly auditable trail of histories, workflows, and datasets. That end-to-end traceability not only reproduces published biological signals and scales to global pathogen discovery, but also resolves unstated sample pooling, annotation mismatches, custom table filtering, and tutorial-subset reference data across the original publications.

## Results

### Advanced models propose vastly different analysis plans …

When handed an RNA-seq dataset, a coding agent returns a plausibly looking workflow in seconds, leaving no easy way to judge whether the plan is sound. We asked 30 independent agents—10 each from Anthropic (Claude Code, Claude Opus 5.5), OpenAI (Codex, GPT-6 Astra), and Google (Antigravity, Gemini 3.8 Flash)—to design a replication plan for a transcriptomic study of the foodborne pathogen *Bacillus cereus* (PRJEB86181), which causes diarrheal illness by secreting pore-forming toxins in the gut [15]. Vyklická et al. tested how 0.5% dietary glucose regulates toxin expression in the dairy-associated strain ATCC 10987 (*n* = 2 glucose vs. *n* = 3 control replicates from a 19-library sugar screen, see Methods). Two pipeline details were left unstated in the methods text: the paper reported locus tags from *Bacillus pacificus* anQ-h4 without naming the reference assembly (while RefSeq had suppressed ATCC 10987 during taxonomic revision), and it did not state whether DESeq2 modeled all 19 libraries together or only the 5 focal samples.

Faced with these ambiguities, agreement among the 30 agents collapsed. Twenty-nine of 30 plans diverged from the published workflow, splintering into 18 distinct eight-step pipelines and 11 incompatible designs across the five choices that alter final gene counts and *p*-values: reference genome, rRNA filtering, aligner, gene annotations, and sample pooling (Figure 1, Supplementary Table S1).

**Figure 1:**
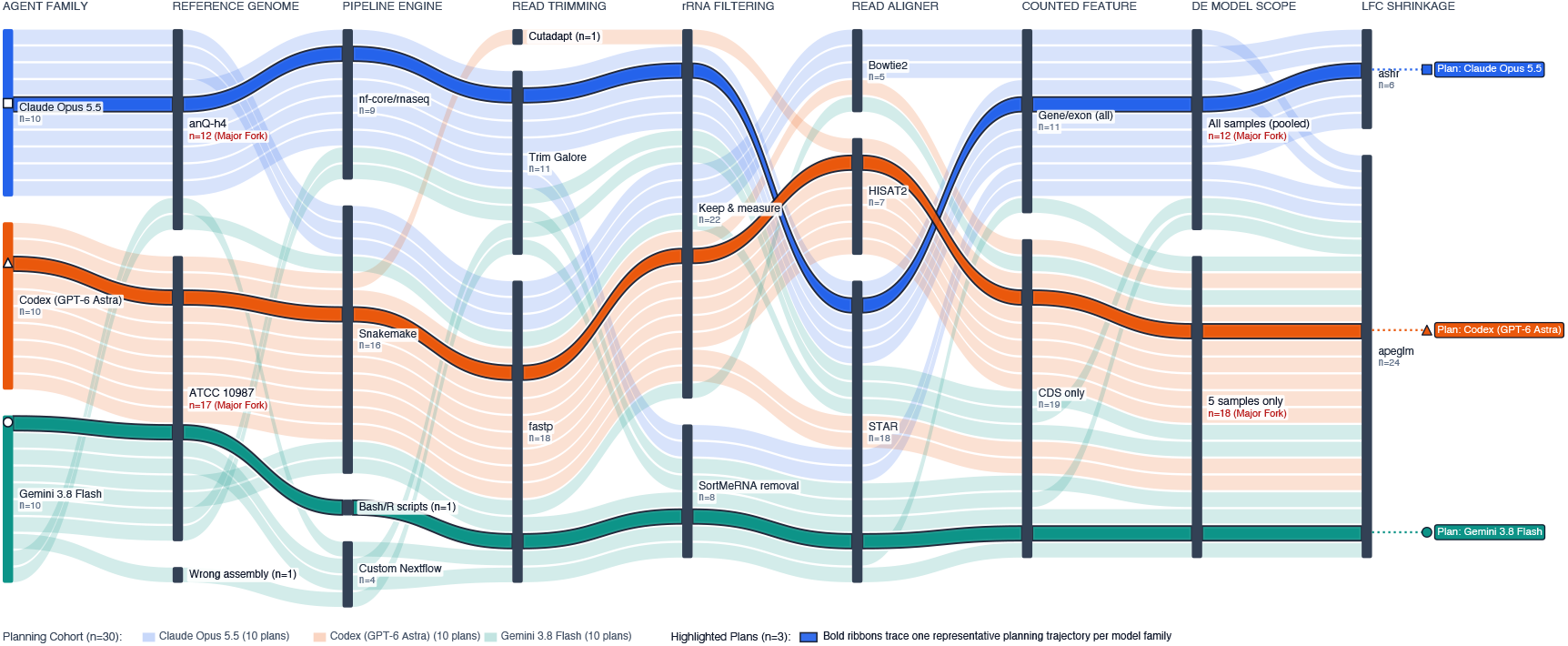
Thirty autonomous agents propose 18 different pipelines for the same bacterial RNA-seq study. Alluvial diagram tracing 30 independent reanalysis plans for a published *Bacillus cereus* carbohydrate-supplementation study (PMC12804731) [15] without access to Galaxy infrastructure. Plans were generated by 10 Anthropic Claude Code agents running Claude Opus 5.5 (blue), 10 OpenAI Codex agents running GPT-6 Astra (orange), and 10 Google Antigravity agents running Gemini 3.8 Flash (teal). Columns represent eight sequential pipeline decisions from reference assembly selection to log-fold-change shrinkage, with node heights proportional to the number of agents choosing each option (*n* = 30 total). Transparent ribbons show individual plans, while bold ribbons with geometric markers trace one executed trajectory per model family (square for Claude, triangle for Codex, and circle for Gemini). Across all eight steps, the 30 plans split into 18 distinct workflows (11 distinct designs across the five result-changing steps), and 29 of 30 differed from the published workflow.

### … and leave a dusty trail of intermediates

We then asked one agent per provider to actually execute its plan locally. Once execution began, each run drifted even further from its own plan as it improvised around local software and archive metadata (Supplementary Table S1). All three runs recovered headline repression of *nheB* and induction of *entFM* (Supplementary Figure S1A), yet diverging reference genomes silently altered the detected toxin repertoire. Aligning to *B. pacificus* anQ-h4 missed cytotoxin K (*cytK2*) entirely because that assembly lacks the locus, whereas aligning to the sequenced isolate *B. cereus* ATCC 10987 recovered active *cytK2* transcription across every library (Supplementary Figure S1B). Compute footprints varied by orders of magnitude, and the three runs left behind 26 unversioned glue scripts totaling over 1,200 lines of non-portable code (Supplementary Tables S1 and S2). Without shared infrastructure, local agents leave a debris field of disposable scripts in place of a verifiable record.

### Agentic science on public infrastructure: The Galactic plane

Eliminating pipeline fragmentation, disposable glue code, and local hardware limits requires pairing domain skills with a shared execution engine (Supplementary Figure S2). Any coding agent—or Orbit, our desktop workbench—connects to public Galaxy infrastructure [8] through modular skills and the Galaxy MCP server [16, 17]. Instead of improvising shell commands, the agent invokes containerized tools and peer-reviewed workflows via typed API calls across national supercomputers and cloud nodes (Supplementary Figure S2, bottom), running everything from memory-intensive aligners like STAR [18] to foundation models and petabyte-scale index searches—such as AlphaGenome [19], Logan [20], and LexicMap [21]—that no personal workstation can host.

Shared infrastructure makes complete provenance mandatory by default (Supplementary Figure S2, top right). Narrative model summaries omit silent parameter changes and failed retries, and few biologists possess version control discipline over interactive shell sessions. Pushing execution onto Galaxy and Orbit records every input file, tool version, parameter, and intermediate dataset in a durable, shareable Galaxy history and workflow graph. The research notebook then links each narrative claim directly to the corresponding Galaxy datasets and job invocations, making the entire analysis human-readable and immediately verifiable. When an analysis needs custom code, the agent registers that script as a versioned, containerized User-Defined Tool^1^ inside the workflow rather than leaving an untracked local file.

### Reproducing a paper using a generic agent on the Galactic plane

To test how shared infrastructure changes interactive analysis, we connected Google’s Antigravity CLI (Gemini 3.8 Flash) to UseGalaxy.org via the Galaxy MCP server [16] to reproduce the *B. cereus* RNA-seq study [15] in a human-in-the-loop session. Unlike local runs that generated 26 unversioned scripts or exhausted laptop disk space, the entire replication ran on free public supercomputers (Supplementary Figure S3).

Shared history and workflow provenance proved essential for human steering. Both the local GPT-6 Astra run and the MCP-connected Gemini 3.8 Flash agent initially ran HISAT2, a eukaryotic spliced aligner that is inefficient for an intronless bacterium (see Methods). Inspecting the shared Galaxy history allowed us to spot the HISAT2 run, steer the agent to assemble a reusable Kallisto workflow [22], and keep the initial HISAT2 output as a direct cross-check (*r* = 0.9622). Quantifying reads against bacterial coding sequences with Kallisto avoids spliced alignment overhead and finishes each library in 38–90 seconds (median 59 seconds) instead of 3.6–6.3 minutes for HISAT2, provided duplicate transcript identifiers on frameshifted transposase features are deduplicated (with both raw and deduplicated references preserved in the Galaxy history, see Methods).

Formal workflow validation also prevented silent scripting workarounds. When the agent wired an incompatible standalone transcript-import step between Kallisto’s output collection and DESeq2— and after GTF-based gene mapping failed at runtime on the bacterial annotation—Galaxy’s workflow editor flagged the collection mismatch in orange so the agent supplied an explicit transcript-to-gene table to DESeq2’s built-in import mode (Supplementary Figure S3A,B). The validated workflow reproduced fivefold repression of *nheB* (*p*_adj_ = 2.99 × 10^−3^), nominally significant nearly threefold repression of *nheA* (*p* = 0.0274, *p*_adj_ = 0.106) and induction of *entFM* (*p* = 0.0158, *p*_adj_ = 0.0730) in the five-library contrast, and unchanged expression of component *nheC* (Supplementary Figure S3D, see Methods). Auditing the resulting Galaxy Page notebook (Supplementary Figure S3C) against the history confirmed that every embedded dataset card and fold-change summary resolves to verified history outputs (Supplementary Table S3).

### An opinionated agent

Domain skills and the Galaxy MCP server guide generic command-line agents effectively, yet terminal agents require command-line fluency and nothing in a skill file stops an agent from falling back on untracked local shell commands when a step fails. To give bench biologists a friendly desktop application whose harness can block and record what the agent does, we built Orbit on top of Pi.dev (Supplementary Figure S2). Orbit is Galaxy-first, steering tool execution to Galaxy while keeping guarded local execution for lightweight work, tracking live cluster jobs, registering custom scripts as versioned User-Defined Tools [23], and syncing the analysis record into a version-controlled project notebook across commercial, public API, and local open-weight models.

Controlling an agent operates across three levels: domain skills that advise how to design an analysis, the MCP server that exposes typed Galaxy tools, and harness logic that mediates every tool call in code. Harness logic is essential because prompt-level instructions can be ignored. Because Galaxy provides a server-side record of typed tool runs that the agent cannot edit, Orbit’s harness logic can verify directly against Galaxy whether a step’s job has actually finished before allowing the plan step to be marked complete (see Methods). These guarantees apply to the record rather than the science: Orbit does not make an agent’s analytical choices correct, but its harness logic ensures that what the agent did is checkable. We used Orbit on UseGalaxy.org to execute all remaining reproductions and discovery analyses summarized in Table 1 and detailed in the sections that follow.

**Table 1:** Summary of empirical validations and discovery analyses executed on the Galactic plane. Across seven biological investigations spanning bacteria, mammalian cortex, fungi, mosquito vectors, protozoan parasites, and 42 viral and bacterial epidemic alignments (41 distinct outbreaks across 37 publications), autonomous agents executed reproducible workflows on free public UseGalaxy.org and ACCESS-CI supercomputing infrastructure. Every analysis produced inspectable Galaxy histories, reusable workflows, and shareable research notebooks (complete accession identifiers in Supplementary Tables S3–S6).

| Analysis & System | Assay & Scale | Agent & Stack | Highlight |
| --- | --- | --- | --- |
| *Bacillus cereus* enterotoxin regulation<br>Vyklícká et al. [15] | Bacterial RNA-seq<br>(5 libraries, 5,472 coding loci) | Generic CLI agent + Galaxy MCP + Skills | Replaces 26 unversioned local glue scripts from unconstrained runs with a validated Galaxy workflow, preserving raw and deduplicated bacterial CDS references in the history and switching from HISAT2 to $\sim 5\times$ faster Kallisto pseudoalignment (38–90 s per library, median 59 s). |
| Mouse cortex spatial transcriptomics<br>Miao et al. [24], Sun et al. [25] | Mammalian scRNA-seq + spatial prediction<br>(3,405 spatial & 5,613 reference cells) | Orbit + 2 UDTs on UseGalaxy.org | Reconciles the 1,000-cell tutorial reference used in Extended Data Fig. 1c against the 5,613-cell Dataset 15 archive cited in Data Availability, reproduces the tutorial subset on Galaxy, and scales to all 5,613 reference cells to triple prediction accuracy ($r = 0.0861$ to $0.2553$). |
| *Aspergillus fumigatus* azole time-course<br>Hokken et al. [26] | Fungal RNA-seq<br>(30 libraries, 3 isolates × 5 time-points) | Orbit + Workflow Foundry + BRC-Analytics + 3 UDTs | Compiles a 60-step workflow from published methods text, shows that the authors fitted one pooled 30-sample model instead of pairwise contrasts, and replaces a decommissioned web service (AspGD) and two manual web steps (Venn overlap and REVIGO) with automated UDTs. |
| *Anopheles gambiae* infection chromatin<br>Ruiz et al. [27] | Vector ATAC-seq<br>(midguts & salivary glands, 4 libraries) | Orbit + 4 UDTs on UseGalaxy.org | Recovers 90.7–93.2% of the 111,586 replicate-shared chromatin regions in the deposited table (81.8–92.2% of all 193,005 deposited peaks), while showing that the paper’s headline counts reflect unmerged per-replicate table rows and tissue-specific overlap filtering. |
| *Aedes aegypti* midgut single-cell atlas<br>Wang et al. [28] | Vector 10x 3’ scRNA-seq<br>(2 replicates, 6,707 post-QC cells) | Orbit + BRC-Analytics on UseGalaxy.org | Stages non-model vector reference data into a public single-cell pipeline, resolves a mitochondrial gene prefix mismatch that breaks generic prefix-based Scanpy QC, and recovers three broad midgut populations and known marker genes. |
| *Plasmodium vivax* VIR phylogenomics<br>Carlton et al. [29], Auburn et al. [30], Minassian et al. [31], Demeulenaere et al. [32], Harrison et al. [33] | Structure-aware phylogenomics<br>(3 genomes, 2,592 proteins) | Orbit + 10 UDTs (1 GPU, 9 CPU) on UseGalaxy.org | Orchestrates GPU protein structure models (ProstT5, Foldseek, and FoldMason) via unprivileged UDTs, collapsing 203 sequence clusters (146 subfamilies with $\geq 3$ members) into 73–76 structural classes (24–30 with $\geq 3$ members) and showing that 20–21 pan-continental classes hold 96.5–97.3% of all VIR proteins. |
| Pathogen emergence dating across 42 cohorts<br>42 BEAST alignments across 37 studies [34–38] | Genomic epidemiology<br>(42 viral & bacterial epidemic alignments, 41 unique) | Orbit + 2 UDTs on UseGalaxy.org | Dates 42 pathogen epidemic alignments in seconds per cohort instead of hours to days of Bayesian MCMC sampling, matching reference Bayesian emergence dates across 61.5–64.1% of scored cohorts. |

### Uncovering a reference discrepancy in spatial transcriptomics study

Because imaging-based spatial transcriptomics measures only a few dozen genes across a tissue section, TISSUE [25] uses paired single-cell RNA-seq reference atlases to impute unmeasured transcripts and calibrate uncertainty intervals. Recently, Miao et al. [24] showcased Paper2Agent by wrapping TISSUE into an agent to predict expression and uncertainty for the vascular smooth muscle marker *Acta2* across 3,405 mouse somatosensory cortex cells (33-gene osmFISH assay). Wrapping code into a standalone agent, however, leaves execution unverified unless input datasets and workflow steps are recorded on shared infrastructure.

Tasked with reproducing the Paper2Agent spatial example, Orbit uncovered an ironic discrepancy: a demonstration designed to showcase automated agent reproducibility had run on a 1,000-cell tutorial subset rather than the full 5,613-cell reference atlas it cited. In Miao et al. [24], human researchers prompted Paper2Agent to follow the TISSUE GitHub tutorial to verify that the agent reproduced a human running the tutorial, while their Methods and Data Availability statement cited the full 5,613-cell mouse cortex reference archive (Zenodo Dataset 15) without noting that the tutorial uses a 96 KB, 1,000-cell subset bundled in the software repository. Because the 1,000-cell tutorial reference omits 82% of the 5,613 reference cells in Dataset 15, it mislocalizes predicted *Acta2* expression into a ventral band and leaves predictions nearly uncorrelated with measured ground truth (*r* = 0.0861, Figure 2A,B,D,E and Methods). Orbit registered two TISSUE User-Defined Tools on UseGalaxy.org, reproduced the 1,000-cell tutorial predictions to floating-point precision (*r* = 1.0000, Δ_max_ *<* 10^−15^, 86.46% Galaxy vs. 86.49% local coverage at nominal 77%, Figure 2A,B), and then ran the same workflow on all 5,613 reference cells in Dataset 15. Using the full reference archive recovered dorsal pial smooth muscle localization, tripled *Acta2* prediction correlation to *r* = 0.2553 (*r* = 0.2845 in 10-fold cross-validation, 85.61% coverage, Figure 2C,F), and improved accuracy across 27 of 33 cortical genes (mean *r* = 0.177 to 0.273, Figure 2G and Supplementary Table S4). Whereas wrapping code into a standalone agent confirms that a tutorial script can be rerun, recording input datasets and workflows in a shared Galaxy history makes explicit what data were actually analyzed.

**Figure 2:**
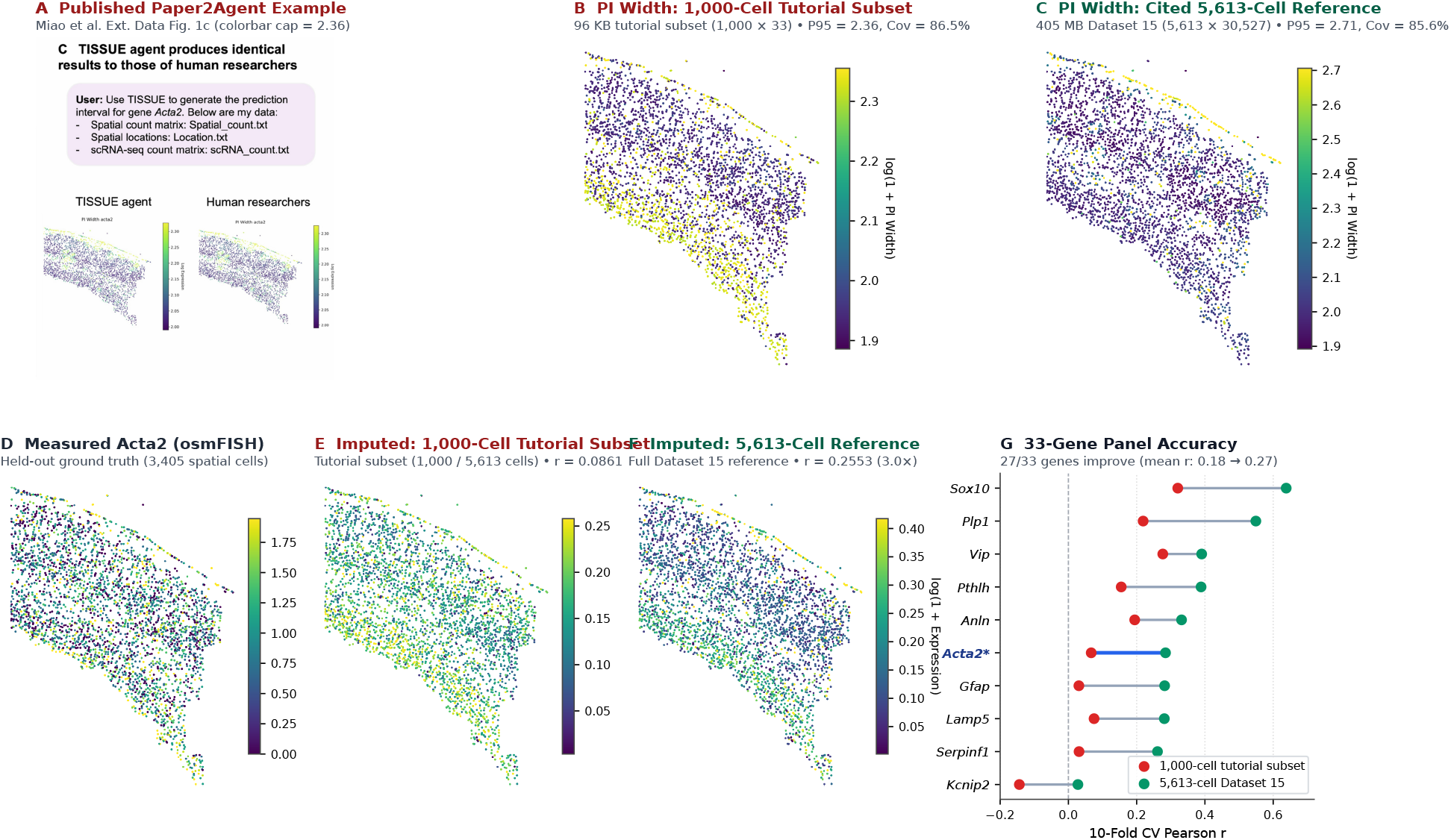
Comparing the 1,000-cell tutorial subset and full 5,613-cell reference atlas for spatial transcriptomics imputation on public infrastructure. **A**, Published Paper2Agent example from Extended Data Figure 1c of Miao et al. [24] showing conformal prediction interval (PI) widths (log(1 + PI Width)) for imputed *Acta2* expression across 3,405 mouse somatosensory cortex cells at nominal 77% coverage (*α* = 0.23). The 95th-percentile colorbar upper bound (≈ 2.36, ticks from 2.0 to 2.3) matches the 96 KB, 1,000-cell tutorial subset rather than the 5,613-cell Dataset 15 reference cited in Data Availability. **B**, Orbit reproduction on UseGalaxy.org using the 1,000-cell tutorial subset (33-gene panel, 31 calibration genes retained after a 1% minimum expression filter drops *Flt1* from 32 unfiltered genes), matching the published 95th-percentile colorbar cap (P95 = 2.3552) and local execution to floating-point precision (*r* = 1.0000, Δ_max_ *<* 10^*−*15^ under identical cross-platform splits, 86.46% Galaxy vs. 86.49% local coverage, mean log(1 + PI Width) of 2.0621 vs. 2.0634 locally). **C**, Scaling the same Galaxy workflow to the cited 405 MB Dataset 15 reference (5,613 reference cells × 30,527 genes, retaining all 32 calibration genes) shifts the 95th-percentile PI width cap to P95 = 2.7062 (mean log(1 + PI Width) = 2.0651, 85.61% coverage) and localizes uncertainty to the pial vasculature. **D**, Measured osmFISH ground-truth expression (log(1 + Counts)) for held-out *Acta2* from Dataset 15, showing true vascular smooth muscle localization along the dorsal pial surface (see Methods). **E**, Imputed *Acta2* expression using the 1,000-cell tutorial subset (*r* = 0.0861), which mislocalizes predicted expression into a diffuse ventral band. **F**, Imputed *Acta2* expression using the full 5,613-cell Dataset 15 reference (*r* = 0.2553, 3.0× higher), recovering the dorsal pial vascular layer. **G**, Leak-free 10-fold cross-validated prediction accuracy (Pearson *r*) across the 33-gene cortical panel comparing the 1,000-cell tutorial subset (red) against the full 5,613-cell Dataset 15 reference (green). Full reference data improves imputation for 27 of 33 genes (mean *r* = 0.177 to 0.273), including *Acta2* (0.067 to 0.285), *Gfap* (0.031 to 0.282), *Serpinf1* (0.031 to 0.261), *Lamp5* (0.075 to 0.281), *Plp1* (0.219 to 0.549), and *Sox10* (0.320 to 0.638).

### Uncovering analysis inconsistencies and defunct tools

To test multi-step functional genomics, we combined Orbit with Workflow Foundry [14]—which compiles published methods prose into validated Galaxy workflows—and BRC-Analytics [39], which stages curated pathogen reference assemblies and sequencing archives (see Methods). We reconstructed the 30-sample RNA-seq study of Hokken et al. [26] in *Aspergillus fumigatus*, an airborne mold causing lethal invasive infections where frontline triazole resistance is rising. Hokken et al. profiled transcriptional responses to itraconazole across three clinical isolates and five exposure timepoints in duplicate (PRJNA482512). BRC-Analytics staged the Af293 genome and all 30 libraries, and Workflow Foundry compiled a 60-step Galaxy workflow (86.88% mean unique alignment vs. 86.5% published, see Methods).

Comparing our initial differential expression outputs against the authors’ supplementary spreadsheet uncovered an unstated statistical modeling choice and three manual downstream web steps. Four separate 12-sample pairwise DESeq2 models reproduced the published fold changes (*r* ≥ 0.998) but inflated standard errors by a median of 20–26% and called 31% and 39% fewer differentially expressed genes at 120 and 240 minutes (missing 34% and 42% of the published DEG sets). Inspecting the supplementary spreadsheet revealed the signature: every gene carried a single identical baseMean value across all four contrasts, showing that the authors had fitted one pooled 30-sample model across all five timepoints simultaneously. With only two replicates per group, pooling all 30 libraries provides 25 residual degrees of freedom to stabilize dispersion and applies a uniform expression threshold (baseMean ≥ 30). Switching our Galaxy workflow to a pooled 30-sample timepoint model matched the published standard error distribution (median ratio 1.000, interquartile range −0.4% to +0.3%) and reproduced DEG counts within 0.3–1.3% at every timepoint (*r* = 0.9996–0.9999, Supplementary Figure S4A), while three User-Defined Tools replaced a decommissioned fungal Gene Ontology server (AspGD) and automated Venn overlap and REVIGO summarization inside Galaxy to recover the shift from cell-wall stress to drug-efflux induction (Supplementary Table S3).

### Uncovering undocumented analysis details

End-to-end provenance also separates reproducible chromatin biology from table-formatting conventions. In the ATAC-seq study of Ruiz et al. [27] across midguts and salivary glands of the malaria vector *Anopheles gambiae*—the two epithelial barriers traversed by *Plasmodium falciparum* parasites during infection (GSE152924)—calling sub-nucleosomal open-chromatin peaks (*<* 130 bp) with the cited MACS2 parameters recovered 90.7–93.2% of the 111,586 replicate-shared regions in the authors’ deposited table (81.8–92.2% across all 193,005 deposited peaks), yet standard consensus peak merging produced less than half of the paper’s headline counts (43,010 midgut and 68,576 salivary-gland regions, which the paper termed unique regions). Comparing the deposited table against our Galaxy outputs showed that the table lists each replicate’s peak as a separate unmerged row and that the stated two-sided 51% overlap rule matches midguts (−3.3%) but under-calls salivary glands (−16.1%, Supplementary Figure S4B, see Methods). Fitting an empirical relaxed overlap threshold reproduced the headline region totals (+4.1% and −1.4%) and matched downstream gene, promoter, and enhancer counts derived from the deposited table within 0.4–2.9% (Supplementary Table S3), preventing unmerged table rows and tissue-specific overlap thresholds from being mistaken for failed peak calling.

### Dealing with disparate annotation formats

Reanalyzing the adult female midgut single-cell RNA-seq atlas of Wang et al. [28] in sugar-fed *Aedes aegypti*—the primary vector of dengue, Zika, chikungunya, and yellow fever viruses—illustrated an annotation trap that affects generic single-cell workflows on non-model organisms. Standard Scanpy pipelines filter dying cells by matching mitochondrial gene prefixes such as `MT-` or `ND`, whereas VectorBase labels *A. aegypti* mitochondrial genes with standard `AAEL` identifiers (which Wang et al. hard-coded as a 38-gene list in their local script). In our Galaxy history, querying a generic prefix matched zero genes and terminated Scanpy quality control, while an empty prefix flagged every gene as mitochondrial. Prefixing the 38 mitochondrial genes in the annotation GTF resolved the failure, retained 6,707 post-QC cells across both replicates (compared with 7,147 published cells after doublet removal), and resolved three broad midgut populations (enterocyte/digestive, hemocyte-like, and visceral muscle) whose sample-blocked pseudobulk profiles recovered known midgut markers and digestive proteases (FDR ≤ 0.002, see Methods and Supplementary Table S3).

### Heavy-Weight Model Orchestration and Structure-Aware Phylogenomics

We next tested whether Orbit can orchestrate GPU protein structure models on free public super-computers to study variant surface antigen diversity in *Plasmodium vivax*, the most geographically widespread human malaria parasite. Each *P. vivax* genome encodes nearly 1,000 VIR variant surface proteins proposed to remodel infected red blood cells and mediate spleen cytoadherence [29–32,40,41]. Because diversifying selection drives VIR sequences into the twilight zone (≤ 30% amino acid identity), sequence clustering across Indonesia (PvP01 [30]), Thailand (PvW1 [31]), and Peru (PvPAM [32]) splits the 2,664 PF05795-positive VIR proteins (2,592 after length filtering) into 203 sequence clusters (146 subfamilies with ≥ 3 members, of which 132 span all three genomes), obscuring family-level relationships across divergent sequences even though structure is conserved in related PIR proteins (where Harrison et al. [33] homology-modeled ten *P. vivax* VIRs onto the *P. chabaudi* CIR fold). Orbit registered ten User-Defined Tools on UseGalaxy.org (one GPU and nine CPU tools) chaining ProstT5, Foldseek 3Di clustering, FoldMason alignment, synteny analysis, and IQ-TREE 3 phylogenetics across ACCESS-CI NVIDIA A100 GPUs and multi-core CPUs (see Methods, Supplementary Table S5).

Structure-aware clustering compressed repertoire diversity across all execution branches, collapsing the 203 sequence clusters (146 subfamilies with ≥ 3 members) into 73 structural classes (24 with ≥ 3 members) on the local CPU reference branch and 76 classes on UseGalaxy.org GPU (25 with ≥ 3 members) and CPU nodes (30 with ≥ 3 members, Supplementary Figure S5A, see Methods). Across every branch, the biological conclusion held: 20 to 21 pan-continental structural classes contained members from all three genomes and accounted for 96.5–97.3% of all VIR proteins (96.8% on Galaxy GPU across 20 classes, 96.5% on Galaxy CPU across 21 classes, and 97.3% locally across 21 classes, or 97.6% of proteins in Galaxy GPU classes with ≥ 3 members), whereas single-genome classes held only 2.0–2.7%. On the 338-tip local reference alignment of the dominant superfamily (FS_001), maximum-likelihood structural phylogenetics and tip-label permutation testing showed no detectable geographic clustering across Indonesia, Thailand, and Peru (parsimony score = 160 vs. null median 160, *p* = 0.55, Supplementary Figure S5B). Although 64.5–65.7% of the 1,672 VIR genes placed on main chromosomes lie within the subtelomeric outer 10% of chromosome arms (57–90% per assembly) and recombination scrambles flanking gene order in all but 11 of 194 local (3 of 141 GPU) reciprocal best-hit pairs, these results are consistent with VIR structural folds diversifying before *P. vivax* spread across continents.

### Dating emergence of key pathogens across 42 benchmark alignments

Finally, we tested high-throughput epidemic dating across 42 viral and bacterial benchmark alignments spanning Ebola, SARS-CoV-2, Zika, Dengue, Chikungunya, Mpox, Influenza, *Mycobacterium abscessus*, and *Yersinia pestis*. We executed this study in Orbit across two separate stages driven by dedicated prompts. First, in a dedicated harvesting and preparation stage, we prompted Orbit to collect, verify, and archive 42 self-contained author BEAST XML files from public repositories and data archives (GitHub, Zenodo, Dryad, and Figshare) across 37 publications (41 outbreaks plus a paired 14.5 kb Ebola coding partition) and unpack their inlined sequence alignments, tip sampling dates, and clock specifications into synchronized Galaxy collections inside a preparation history on UseGalaxy.org (see Methods). Second, because estimating the time to the most recent common ancestor (*t*_MRCA_) from these configurations in BEAST [42] typically requires hours to days of Bayesian MCMC sampling per alignment, a companion analysis prompt directed Orbit to register a User-Defined Tool wrapping ChronAeon (https://github.com/veg/chronaeon) [38] on UseGalaxy.org and date all 42 alignments concurrently in seconds of manifold computation per dataset inside an analysis history (Figure 3, see Methods).

**Figure 3:**
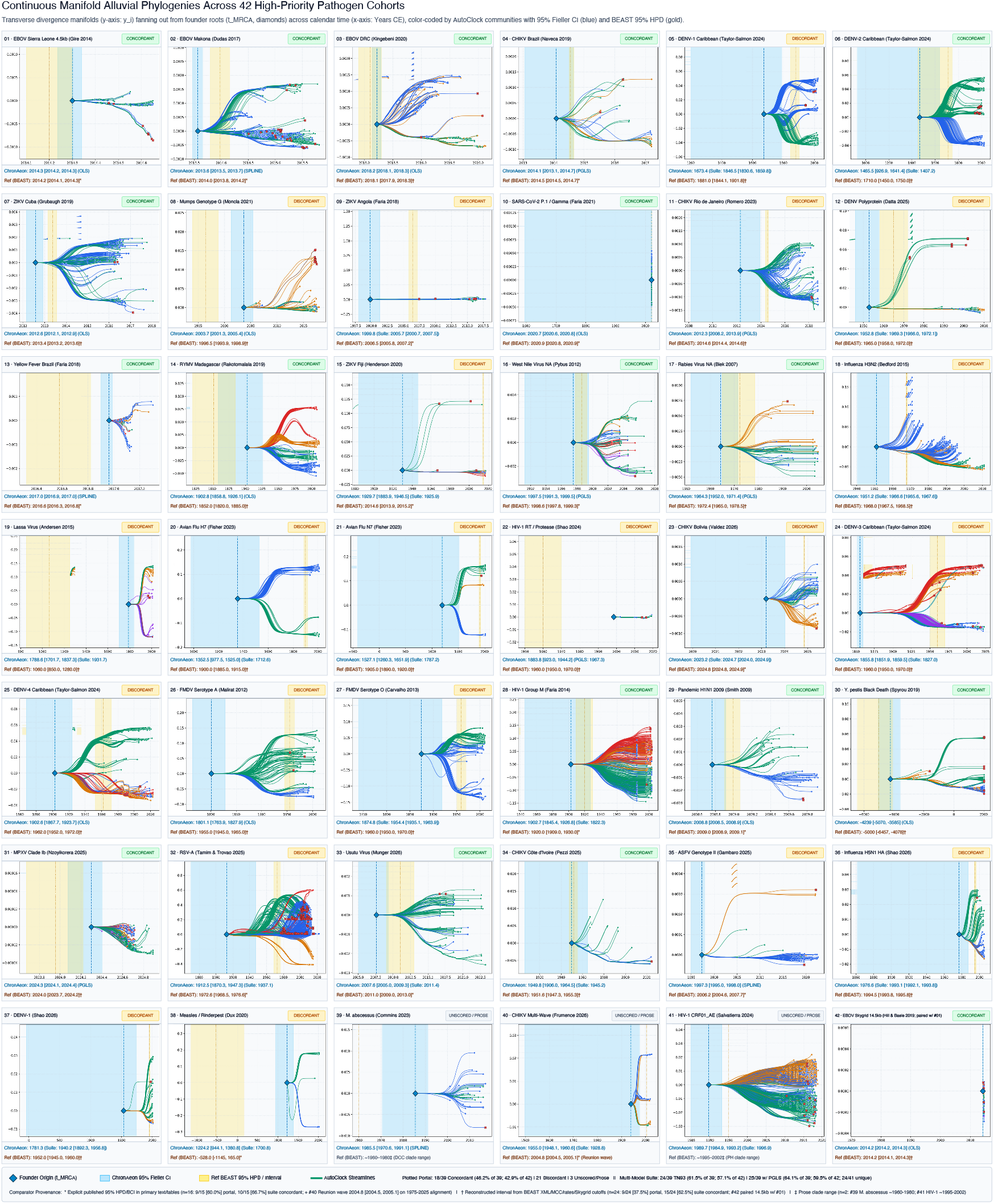
Continuous manifold alluvial phylogenies and emergence dating across 42 pathogen cohorts on UseGalaxy.org. Each panel displays continuous manifold trajectories fanning out from the inferred epidemic origin (*t*_MRCA_, blue diamond) across calendar time, color-coded by AutoClock evolutionary communities (*k* ∈ [0, *K*^*\**^ − 1]) to show lineage expansion and replacement. Shaded bands compare the ChronAeon 95% Fieller analytical confidence interval (light blue) against the published or reconstructed Bayesian reference interval (gold). Top banners and bottom status strips report the pathogen cohort, primary citation, and concordance of the plotted single-pass baseline against the 39 numeric reference intervals (interval overlap or |Δ*t*_MRCA_| *<* 1.0 year). Pairwise-distance regression and HyphAeon Attention PGLS match reference Bayesian dates on 25 of 42 alignments (64.1% of the 39 scored comparators, Supplementary Table S6).

Across major recent outbreaks—including 2014 West African Ebola [34–36] and SARS-CoV-2 Gamma [37], where ChronAeon’s estimate (2020.71) fell within two months of the published BEAST root (2020.87)—tree-free pairwise-distance regression matched reference dates across 61.5% of the 39 scored cohorts (24 of 39), rising to 64.1% (25 of 39, median error 0.40 years across 19 post-1980 outbreaks) with HyphAeon Attention PGLS on Mpox Clade Ib (Supplementary Table S6). Preserving every alignment and diagnostic output in a public Galaxy collection also explained the 14 discordant cohorts: unrooted distance regression breaks down on ancient roots compressed by purifying selection and mutational saturation (such as Lassa [43, 44] and Measles/Rinderpest [45]) and on multi-lineage alignments where whole-alignment regression dates a deep species split rather than a regional introduction (see Methods).

### Putting this all together

Anyone can use this architecture today (https://galaxyproject.org/agents/) with a free account and API key on UseGalaxy.org, UseGalaxy.eu, or UseGalaxy.org.au. Command-line users can install agentic-plugins (https://galaxyproject.org/agents/plugins/) to register galaxy-mcp with curated Galaxy and Workflow Foundry skills, while desktop users can download Orbit for macOS, Linux, or Windows (https://galaxyproject.org/agents/orbit/) using commercial APIs, Chat-GPT subscriptions, or free local Ollama models. Both paths execute every step as a verifiable Galaxy job, and we are embedding a native open-model agent directly inside the Galaxy web interface.

## Discussion

Autonomous agents make complex data analysis feel effortless, allowing any student or investigator to generate polished figures in minutes. As our benchmark and reanalyses show, however, effortless execution routinely masks consequential choices: surrogate reference assemblies from a related strain that lack expressed toxin genes, inefficient eukaryotic spliced aligners applied to bacteria, unpooled dispersion models, or reversed library strandedness. Equally instructive is what our reproduction of the Paper2Agent spatial transcriptomics example [24] revealed. To verify that an agent could follow the TISSUE software tutorial, Miao et al. prompted Paper2Agent with the repository’s 1,000-cell tutorial files while citing the full 5,613-cell mouse cortex reference atlas [25] in their Methods and Data Availability statement, leaving a threefold gap in prediction correlation between the tutorial subset analyzed in the figure and the full reference dataset cited in the text (*r* = 0.0861 vs. *r* = 0.2553, see Methods). Wrapping code into an agent in a local environment shows that a tutorial command can be rerun, but it does not record which datasets entered the computation. Connecting agents to public Galaxy infrastructure, visual workflow validation, and national supercomputers makes every input dataset, parameter setting, and analytical step immediately inspectable by authors, collaborators, and reviewers.

Several practical limitations clarify where human expertise and community guardrails remain essential. Whereas our planning benchmark evaluated *n* = 10 independent runs per model family, our live execution comparisons and biological case studies represent interactive sessions that incorporated investigator steering—such as switching from an initial HISAT2 run to a Kallisto workflow in *B. cereus*—alongside agent-driven reanalyses. Connecting agents to public infrastructure also does not eliminate model-level execution failures, and the record is what makes them diagnosable—failures, including defects in Orbit itself, can be identified after the fact from each session’s notebook and activity log rather than from the agent’s own summary. At the same time, while Galaxy records complete server-side tool, container, and dataset lineage, agent-level metadata—such as which language model, thinking-effort setting, or multi-model switch submitted each job—is currently recorded only in Orbit’s local session logs rather than stamped onto Galaxy job tags, highlighting a clear next step for server-side provenance integration. Shared infrastructure likewise does not eliminate minor floating-point or clustering differences across software environments, the ChronAeon clock module was evaluated by its developers (with 24 of 39 reference intervals reconstructed from prose or clock calibrations, see Methods), and commercial model APIs still carry usage costs. In our evaluations, however, skill-guided workflows perform well with open-weight models such as DeepSeek at a fraction of the cost, and serving open models directly on public supercomputers will remove that barrier altogether.

Running agents on public infrastructure also requires balancing extensibility against security and software sprawl. Historically, Galaxy users could only run pre-installed tools, which stalls an agent whenever an analysis needs a custom script or new model. User-Defined Tools (UDTs) solve this by packaging custom scripts or heavy models into isolated containers on demand, but arbitrary container execution on shared clusters requires access controls: UseGalaxy.org grants UDT privileges via an approved account list, whereas UseGalaxy.eu routes requests through an automated packaging and pull-request review pipeline. As binary-decompilation agents such as Reverse Engineer Anything (REA) [46] make software replaceable on demand, thousands of uncurated UDTs could easily degenerate into an unmaintained graveyard no better than thousands of abandoned GitHub repositories, where stars reflect popularity rather than whether code still runs on real data. Shared execution on Galaxy provides a concrete selection and governance mechanism across three stages (see Methods). First, agents register UDTs with mandatory pinned containers and typed schemas in user-scoped histories to unblock immediate analyses without cluttering the global tool panel. Second, platform execution telemetry tracks real-world invocation counts, failure rates, and resource footprints across independent users while flagging duplicate wrappers. Third, recurring high-utility UDTs are routed to the Intergalactic Utilities Commission (IUC) [10] and Intergalactic Workflow Commission (IWC) [11], where human maintainers consolidate redundant wrappers, add Planemo [47] regression tests, and graduate vetted tools into permanent community standards.

Finally, verifiable agentic infrastructure changes what a scientific publication itself represents. When an agent executes a study on public infrastructure, the primary scientific product is no longer a static prose manuscript—it is the immutable Research Object Crate (RO-Crate) of inputs, workflows, container digests, and results archived on Zenodo with a permanent DOI. Once that computational lineage is locked and auditable, multimodal synthesis tools such as NotebookLM can generate written reports, slide decks, or interactive summaries on demand in whatever format a reader prefers, letting scientific communication adapt to any medium without sacrificing rigor.

## Methods

### Multi-Agent Analytical Divergence Benchmark

We evaluated analytical divergence across autonomous agents using an RNA-seq study of carbohydrate-mediated enterotoxin repression in *Bacillus cereus* (BioProject PRJEB86181, PMC12804731) [15]. The primary contrast compared two 0.5% (w/v) glucose-supplemented replicates (ERR14660412 and ERR14660413) against three baseline tryptic soy broth control replicates (ERR14660420, ERR14660421, and ERR14660422) from nineteen RNA-seq libraries across seven growth conditions (six carbohydrate supplements plus tryptic soy broth control, alongside one whole-genome sequencing run, ERR14660405, totaling twenty ENA runs, where the authors’ Supplement S4 reports a single identical `baseMean` per gene across all six contrasts from a pooled 19-library DESeq2 model). In the published study, excess glucose represses the primary pore-forming enterotoxin subunit *nheB*, leaves component *nheC* unchanged, and induces the cell-wall peptidase *entFM*, while diarrheal toxins hemolysin BL (*hbl*) and cytotoxin K (*cytK*) are reported as unexpressed.

To quantify planning variation, we evaluated 30 isolated agents across three foundation model families: ten Claude Opus 5.5 agents (Anthropic Claude Code CLI, cc), ten Codex agents running GPT-6 Astra (OpenAI Codex CLI, `codex exec`), and ten Gemini 3.8 Flash agents within Antigravity (`agy`, Google). The version strings designate the frontier Anthropic, OpenAI, and Google CLI model snapshots evaluated in our early-2026 benchmark logs. Each agent received identical natural-language instructions in an empty local workspace directory with no prior session context, external memory, or access to Galaxy cyberinfrastructure (Table 2). During planning, each agent operated under its native CLI tool permissions: web fetch/search and curl in Claude Code, live web search in a network-restricted shell sandbox in Codex, and URL reading plus shell execution in Antigravity. We parsed each generated plan across eight sequential analytical decisions: reference genome assembly, workflow engine, read trimming software, ribosomal RNA depletion strategy, read aligner, quantified genomic feature, differential expression model scope (19-sample pooled vs. 5-sample contrast-only), and log-fold-change shrinkage prior. Combinatorial trajectory diagrams were constructed from this discrete decision matrix. Model families exhibited distinct planning profiles shaped by both model priors and CLI tool permissions. Claude plans aligned to *Bacillus pacificus* anQ-h4 and pooled variance across all nineteen RNA-seq libraries. Codex plans aligned to the sequenced isolate *Bacillus cereus* ATCC 10987 and restricted the comparison to the five target replicates. Gemini plans showed severe instability across assemblies, citing a non-*Bacillus* assembly (`GCF_000008105.1`, *Salmonella enterica* serovar Choleraesuis SC-B67) mislabeled as *B. cereus* ATCC 10987 in one plan, citing the *B. cereus* ATCC 14579 chromosome accession (`AE016877.1`) for ATCC 10987 in another, and reversing library strandedness in four plans. Aligning against a wrong assembly or genus alters or invalidates gene coordinates and annotations. Inverting library strandedness discards or miscounts real transcripts. Pooling all nineteen libraries to estimate variance alters statistical power compared to analyzing only the five target replicates.

**Table 2:**
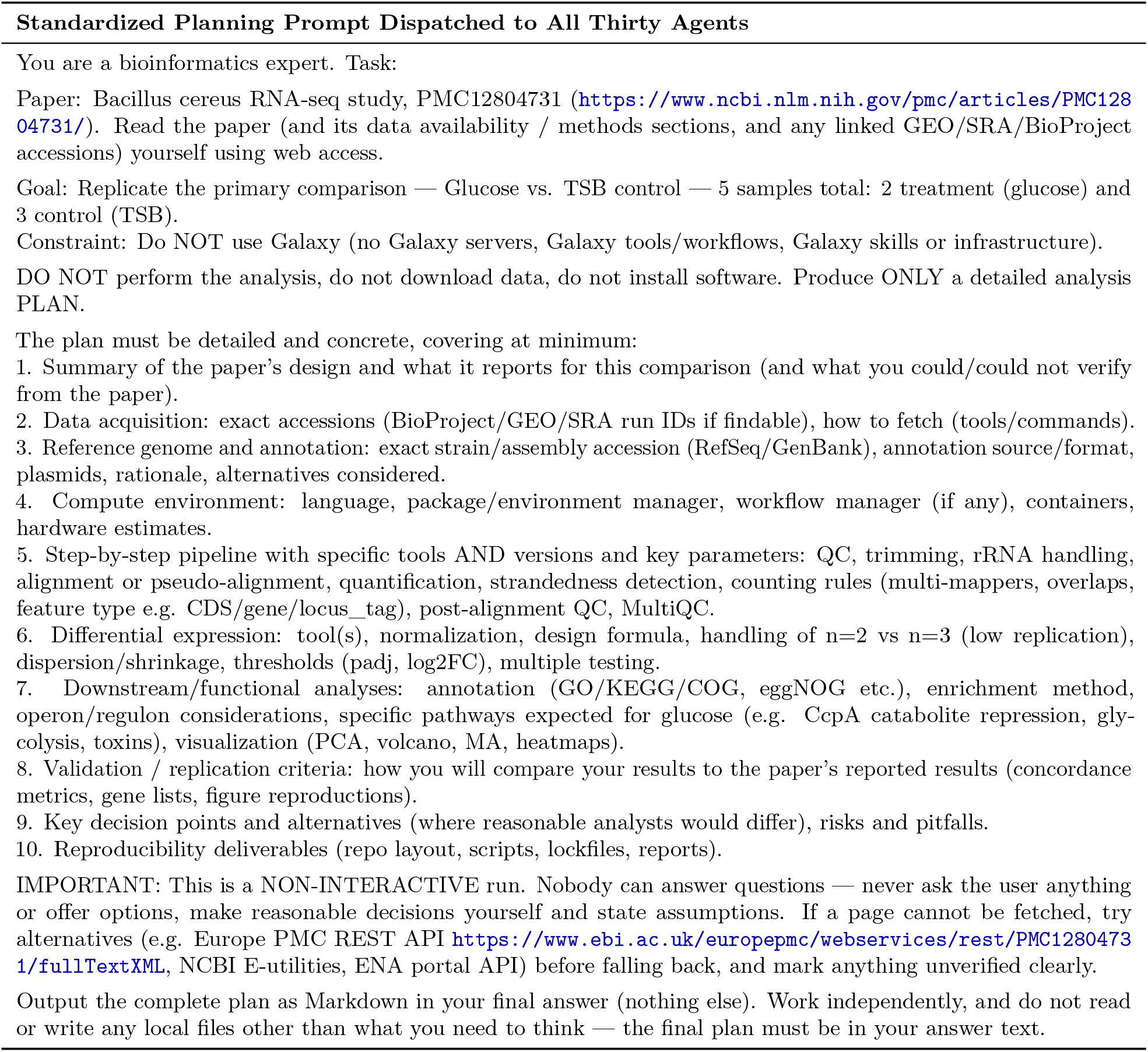
Standardized natural-language prompt dispatched to thirty autonomous agents. All agents received this identical prompt in an isolated workspace with no inter-session memory or access to Galaxy cyberinfrastructure. The prompt directed agents to formulate an end-to-end analysis plan across ten explicit criteria without human intervention or external workflow platforms.

For empirical re-analysis, three agents executed end-to-end replications on dedicated host hardware. Runs used Gemini 3.8 Flash (Antigravity CLI), Claude Opus 5.5 (Claude Code), and Codex (GPT-6 Astra) on an Ubuntu Linux workstation (24 CPU cores, 62 GB RAM, NVMe storage). During live execution, each agent was instructed to fetch the five target RNA-seq libraries from the European Nucleotide Archive, perform quality control, align reads, quantify gene-level abundances, and estimate differential expression for target enterotoxin genes (*nheA, nheB, nheC, entFM, cytK*, and the *hbl* operon), with freedom to select its alignment and modeling stack. Gemini executed an ad-hoc bash script driving Bowtie2 v2.5.5 and unstranded featureCounts v2.1.1 on *Bacillus pacificus* anQ-h4 (unshrunken Wald log_2_ FC = −2.41, *p*_adj_ = 0.0021 for *nheB*, and log_2_ FC = +1.42, *p*_adj_ = 0.0658 for *entFM*, with *nheC p*_adj_ = NA after independent filtering, raw *p* = 0.51). Whereas all ten Claude planning runs proposed pooling all nineteen libraries on *B. pacificus* anQ-h4, during live execution Claude Code analyzed the five target samples and aligned reads against the sequenced isolate *Bacillus cereus* ATCC 10987 (`GCF_000008005.1` / `GCA_000008005.1`, NCBI BioProject `PRJNA74`, where RefSeq later suppressed `GCF_000008005.1` due to an unverified source organism during the *B. cereus*/*B. pacificus* taxonomic revision). The Claude Code run used containerized nf-core/rnaseq 3.14.0 and nf-core/differentialabundance 1.4.0 under Docker, consuming over 100 GB of scratch disk. Aligning to ATCC 10987 recovered active baseline transcription of the cytotoxin K locus (`BCE_1209` / `BCE_RS06135`, 548–1,420 raw read pairs per library, mean 797, DESeq2 baseMean = 831, non-differentially expressed, log_2_ FC = +0.58, *p*_adj_ = 0.43), which was reported as undetected in the published study, consistent with alignment against *B. pacificus* anQ-h4 (an assembly lacking *cytK*). Claude Code’s nf-core/differentialabundance pipeline output `ashr`-shrunken posterior effect sizes and standard deviations, replicating *nheB* repression (log_2_ FC = −2.08, *p*_adj_ = 0.0002) and *entFM* directional induction (log_2_ FC = +1.08, *p*_adj_ = 0.0620). Codex replaced its planned Snakemake pipeline with fourteen custom Bash, Python, and R scripts executing HISAT2 and unshrunken DESeq2 Wald tests with dual-reference sensitivity checks across both anQ-h4 and ATCC 10987 (replicating *nheB* log_2_ FC = −2.16 to −2.46, *p*_adj_ *<* 10^−4^, and *entFM* log_2_ FC = +1.32 to +1.35, *p*_adj_ ≤ 0.0220, with anQ-h4 *nheC p*_adj_ = `NA`, raw *p* = 0.66). We captured all generated shell scripts, Conda environments, Docker layer footprints, and execution logs to measure toolchain overlap, disk consumption, and provenance persistence (Supplementary Tables S1 and S2).

### Galaxy-Native Replication of *Bacillus cereus* RNA-seq

To evaluate interactive agent-assisted replication on public cyberinfrastructure, we connected the Antigravity CLI (agy) to UseGalaxy.org through the Galaxy MCP server (`galaxy-mcp`) [16]. The five target RNA-seq libraries from BioProject PRJEB86181—two 0.5% glucose-supplemented replicates (ERR14660412 and ERR14660413) and three tryptic soy broth control replicates (ERR14660420, ERR14660421, and ERR14660422)—were imported into a Galaxy history alongside the *Bacillus cereus* ATCC 10987 reference genome (`GCF_000008005.1` / `GCA_000008005.1`, BioProject `PRJNA74`) and coding sequence transcriptome (4.59 Mb across 5,476 FASTA records mapping to 5,472 unique coding loci, Supplementary Table S3).

The agent initially executed a HISAT2 and featureCounts pipeline across the five libraries on the full 5.43 Mb genome assembly (HIDs 1–64, cloned into companion history `bbd44e69cb8906b5d979` `c5818e07f6ad`). Inspecting the shared history allowed us to spot the eukaryotic spliced aligner and direct the agent to assemble a prokaryotic Kallisto pseudoalignment workflow while retaining the HISAT2 run as a cross-check. Extracting coding sequences from `GCF_000008005.1` with `gffread` produced four duplicate `unassigned_transcript_N` identifiers across split frameshifted transposase features (`BCE_0021`, `BCE_2178`, `BCE_3634`, and `BCE_4277`). Because Kallisto index construction rejects duplicate target identifiers, the agent deduplicated these headers locally and uploaded the deduplicated CDS reference (HID 106) alongside the raw CDS export (HID 66), preserving both in the public history after verifying the step on a 1,000-read pilot subset.

The agent then assembled and validated a reusable four-step Galaxy workflow combining `fastp`, `kallisto` `quant` [22], `deseq2`, and `multiqc` (Supplementary Table S3). During workflow construction and testing, Galaxy’s workflow schema validator flagged a standalone `tximport` step wired between Kallisto’s output collection and DESeq2’s collection-mode input in orange (Supplementary Figure S3A), and runtime GTF-based gene mapping failed on the prokaryotic annotation (HIDs 144 and 146). The agent therefore removed the standalone `tximport` step, uploaded a two-column transcript-to-gene mapping table (`tx2gene.tsv`, HID 148), and configured DESeq2’s integrated `tximport` mode to aggregate collection-level transcript abundances into gene-level counts (Supplementary Figure S3B). Under glucose supplementation in this five-library single-contrast design (*n* = 2 glucose vs. *n* = 3 control replicates), primary lytic enterotoxin subunit *nheB* decreased 5.05-fold (log_2_ FC = −2.335, *p*_adj_ = 2.99 × 10^−3^). Subunit *nheA* decreased 2.80-fold (log_2_ FC = −1.484, raw *p* = 0.0274, *p*_adj_ = 0.106, HISAT2 *p*_adj_ = 0.0917), cell-wall enterotoxin *entFM* was induced 2.70-fold (log_2_ FC = +1.432, raw *p* = 0.0158, *p*_adj_ = 0.0730, HISAT2 *p*_adj_ = 0.0785), and component *nheC* remained unaffected (log_2_ FC = −0.158, raw *p* = 0.843, *p*_adj_ = 0.907). Both *nheA* and *entFM* reached nominal significance (raw *p <* 0.05) in the five-library model while falling just above the 5% false discovery rate threshold after Benjamini–Hochberg adjustment across the 2,815 loci passing DESeq2 independent filtering (compared with Vyklická et al.’s 19-library pooled model). On UseGalaxy.org, Kallisto pseudoalignment completed in 38–90 seconds per library across the two workflow invocations (median 59 seconds, 38–60 seconds in the primary invocation) without generating intermediate BAM files, compared with 3.6–6.3 minutes per library for HISAT2 alignment and coordinate sorting on standard Galaxy workers (∼ 5× faster, and 4.0–4.8 hours when HISAT2 jobs encountered worker contention, Supplementary Table S3). Across the 5,250 expressed bacterial coding loci with non-missing fold-change estimates (of 5,472 total coding loci), Kallisto pseudoalignment and HISAT2 spliced alignment produced near-identical log_2_ fold-change estimates (Pearson *r* = 0.9622, *R*^2^ = 0.9257). Auditing the interactive Galaxy Page notebook against the replication history confirmed that all embedded dataset references and fold-change tables resolve to verified history outputs (Supplementary Table S3).

### Galaxy MCP Server and Modular Skills

Agents interact with computational infrastructure through the Galaxy MCP server (`galaxy-mcp`) [16], which implements an open standard for tool execution and data inspection. Instead of composing shell commands or ad-hoc API scripts for Galaxy work, agents call the server’s structured operations (45 in `galaxy-mcp` v1.11.2). These endpoints enable agents to search tool registries, inspect dataset collections, launch workflows, and monitor execution states through standard input-output streams or remote HTTP connections. The reference server is implemented in Python using the BioBlend library to communicate with the Galaxy REST API. A TypeScript implementation of the same operation set (@galaxyproject/galaxy-ops) is held to a shared contract with the Python server, and Orbit v0.8.0 exposes the Python server to its agent. Transport mechanisms include standard input-output (stdio) for local CLI integration and streamable HTTP for web and remote clients. Authentication occurs through long-lived Galaxy API keys or OAuth2 token exchanges supplied via environment variables (`GALAXY_API_KEY`). The server provides structured error handling and schema validation across all parameter endpoints.

Domain expertise is provided through modular skill packages [17] that teach language models standard bioinformatics conventions. Skills guide tool discovery, parameter selection, and collection manipulation idioms. For complex multi-step pipelines, Workflow Foundry skills compile validated workflow templates into agent memory [14]. By separating operational skills from the underlying compute runtime, the framework steers agents toward community-vetted analytical procedures rather than improvised ones. Skills are guidance, not enforcement—an agent can still deviate from them, which is why the execution record matters.

### User-Defined Tool Architecture

User-Defined Tools are written in YAML adhering to the `GalaxyUserTool` class definition. The tool wrapper specifies a mandatory container image hosted on public registries (Quay.io, Docker Hub), typed input parameters, declared output datasets, and an execution command. Commands are templated using sandboxed ECMAScript evaluation executed inside an isolated V8 runtime. Output files are claimed from the working directory through `from_work_dir` designations or dynamic dataset discovery patterns. On public servers, UDT execution is governed by instance-specific access controls (approved account access lists on UseGalaxy.org and automated pull-request packaging queues on UseGalaxy.eu) and platform execution telemetry that tracks invocation counts, failure rates, and resource usage to route recurring wrappers to the Intergalactic Utilities Commission (IUC) [10] and Intergalactic Workflow Commission (IWC) [11] for Planemo [47] regression testing.

### Orbit Workbench and Execution Guard

Orbit is an Electron desktop application around a headless agent runtime built as an extension on Pi.dev, and the same runtime also runs as a terminal client. The description below reflects Orbit v0.8.0. The analysis directory is the unit of work. Plans, decisions, and results are Markdown sections of a single `notebook.md`, which Orbit versions automatically in Git repositories it manages. When the agent submits a Galaxy workflow or tool run, it records the run in the notebook as a YAML block (`loom-invocation`, `loom-job`). From then on, a background poller checks Galaxy every 15 seconds without invoking the model, writes the run’s status back into the block (text the model can still edit in this release), and flags a workflow that has failing jobs while others are still running. Galaxy finishing a run is kept separate from the result being checked. When a run completes or fails, Orbit prompts the agent to verify the outputs or investigate the failure so the researcher does not have to relay the notification (sending at most three consecutive follow-ups by default, with user input or Stop taking precedence). The notebook can be bound to a Galaxy Page on the analysis history and pushed to it, storing the narrative record alongside the datasets it describes. Orbit’s local session logs (`session.jsonl` and `activity.jsonl`) also record which foundation model, provider, and thinking-effort level executed each conversational turn (including multi-model switching within a project, such as `gpt-5.5`, `claude-opus-4-7`, `gemini-3.1-pro-preview`, `gemini-3.5-flash`, and `deepseek-v4-pro` across the *P. vivax* phylogenomics session), whereas server-side Galaxy job records currently store tool, container, and dataset provenance without the upstream language-model identifier.

Orbit enforces some rules in code and advises others under its default permission settings. An execution guard intercepts `bash`, `read`, `write`, and edit calls through Pi.dev lifecycle hooks and returns allow, ask, or deny from classifiers for shell hazard, sensitive-file reads, path containment, and model tier, where models classified as weaker by name and price receive stricter treatment of sensitive and out-of-workspace file access. Writes outside the workspace and its configured roots require approval, reads of recognized credential stores are denied, and the API keys Orbit holds are redacted by value from tool results and from the result summaries in the session activity log (`activity.jsonl`). Recognized destructive Galaxy operations require user confirmation and are denied when no user is present to give it. Duplicate or fanned-out SRA imports through the standard download tools are refused before they reach Galaxy, and /`execute` and /`run` refuse to start without a readable notebook or, for a plan with pending remote steps, a Galaxy connection. An optional operating-system sandbox confines shell writes beneath the guard. Skills are advisory, as are two checks on the plan: when a plan is drafted without consulting the IWC workflow registry, Orbit sends one follow-up asking the agent to check, and in v0.8.0 it logs, without blocking, a plan step marked complete while the workflow invocation bound to it is still in progress. In our harness evaluations on open-weight models served through TACC (`evals`/ in the Orbit repository), a prompt-only rule requiring approval before writing a plan to the notebook was followed in 0 of 15 graded runs (Qwen3 collapsed the four-stage protocol to two stages, and Llama 3.3 wrote to the notebook on its first turn). In response, the development version following v0.8.0 refuses by default to let a plan step be marked complete while its bound Galaxy run is still in progress—a default adopted after twelve scripted sessions, each run in triplicate, showed that every refusal was a true positive and that honest completions recorded in notebook blocks were let through (`docs/evidence-gate-audit.md`).

### Workflow Foundry and BRC-Analytics

Galaxy Workflow Foundry (@galaxy-foundry/gxwf-foundry) [14] is a skill-compilation pipeline implemented in TypeScript and Node.js that converts reusable procedural templates (Molds) into agent skills for diverse agent runtimes such as Claude Code, Codex, Antigravity, and Orbit. Each Mold is a Markdown specification carrying Zod-validated YAML frontmatter that declares upstream handoff schemas, design patterns, reference manuals, and downstream output artifacts. Instead of prompting an agent to invent a multi-step workflow in a single pass, Foundry decomposes workflow synthesis into staged skills that pass structured JSON and Markdown handoffs. Staged compilation extracts a structured summary from a published paper, Nextflow pipeline, or CWL workflow, designs the Galaxy interface and collection data-flow topology, resolves curated ToolShed wrappers or User-Defined Tools, and assembles a `gxformat2` workflow step by step. Synthesized workflows are validated statically using the `gxwf` validation engine, which verifies graph topology, step connectivity, and tool parameter schemas against official Galaxy XML definitions and Intergalactic Workflow Commission (IWC) exemplars [11] before automated execution testing via Planemo [47].

BRC-Analytics (https://brc-analytics.org) [39] is the NIAID Bioinformatics Resource Center analytics portal built on public Galaxy infrastructure. It maintains a taxonomy-indexed catalog of curated pathogen, fungal, eukaryotic parasite, and vector reference assemblies, gene annotations, and public sequencing runs aggregated from NCBI Assembly, RefSeq, Ensembl, VEuPathDB, and the Sequence Read Archive (SRA/ENA). In autonomous workflows, BRC-Analytics removes the error-prone step of scraping FTP directories, reconciling assembly accession versions across databases, or downloading raw sequencing files to local storage. Agents query BRC-Analytics by taxonomic identifier or BioProject accession to resolve canonical reference FASTA sequences, GTF/GFF3 gene models, and paired or single-end SRA run collections and stage them into a target UseGalaxy.org history with preserved sample metadata.

### Spatial Transcriptomics Conformal Uncertainty Reproduction

To reproduce and scale the spatial transcriptomics example of Miao et al. [24] and Sun et al. [25], Orbit registered User-Defined Tools `tissue_spatial_prediction` v1.2.0 and `tissue_full_evaluation` v1.0.0 (container quay.io/biocontainers/squidpy:1.5.0, 4 CPU cores) on UseGalaxy.org (Supplementary Table S4). The tools implement SpaGE principal-vector domain adaptation (10 principal vectors) and fixed-radius *k*-nearest-neighbor spatial graph construction (*k* = 15). They then apply cell-centric variability stratified conformal prediction (*k*_*g*_ = 4 gene groups, *k*_*c*_ = 2 cell groups, 10-fold cross-validation, random seed 42 for single-gene *Acta2* and 444 for 33-gene evaluation) at nominal coverage 1 − *α* = 0.77 (*α* = 0.23) and 0.80 (*α* = 0.20). Single-gene *Acta2* prediction and 10-fold spatial cross-validation workflows across all 33 osmFISH panel genes were executed on UseGalaxy.org across both the 1,000-cell repository tutorial subset and the complete 5,613-cell Zenodo Dataset 15 reference archive.

### Reconciling Tutorial and Full-Reference Datasets in the Paper2Agent Spatial Example

In their Methods and Data Availability statement for the mouse somatosensory cortex osmFISH spatial prediction example (Extended Data Figure 1c), Miao et al. [24] cited the Zenodo Dataset 15 archive of Sun et al. [25] (Zenodo record `8259942`), which contains 3,405 osmFISH spatial cells profiled across 33 genes paired with a 5,613-cell single-cell RNA-seq reference atlas profiled across 30,527 genes. In their Supplementary Note and Methods, Miao et al. record that human researchers instructed Paper2Agent to follow the TISSUE GitHub tutorial using a natural-language prompt that named the repository tutorial files (`tests/data/Spatial_count.txt`, `tests/data/Locations.txt`, and `tests/data/scRNA_count.txt`) to verify that the agent reproduced a human running the tutorial. Inspecting those repository files showed that `tests/data/scRNA_count.txt` is a 96 KB, 1,000-cell tutorial and continuous-integration subset (1,000 reference cells × 33 osmFISH panel genes, with tests/data/Spatial_count.txt pre-filtered to 3, 405 × 32 genes after dropping held-out *Acta2*), omitting 4,613 of the 5,613 reference cells (82.2%) in the cited Zenodo Dataset 15 archive. On this 1,000-cell tutorial subset (31 calibration genes after TISSUE’s default 1% *Flt1* filter, or 32 unfiltered), local and Galaxy UDT runs matched predicted *Acta2* expression to floating-point precision (*r* = 1.0000, Δ_max_ *<* 10^−15^), but correlation with measured ground-truth *Acta2* expression from Dataset 15 across the 3,405 spatial cells was only *r* = 0.0861. Stochastic *k*-means stratification on the 1,000-cell subset yielded empirical coverage of 86.46% on Galaxy (86.49% locally) at *α* = 0.23, mean log(1 + PI Width) of 2.0621 (2.0634 locally), and colorbar range [1.89, 2.35] (95th percentile 2.3552), matching the [2.00, 2.32] range plotted in Extended Data Figure 1c of Miao et al. (where *k*-means stratification shifts individual high-uncertainty cells between the ventral boundary and pial edge). When we executed the identical Galaxy workflow on the full 5,613-cell Dataset 15 reference archive (subsetted to the 33 shared panel genes), single-gene *Acta2* prediction correlation tripled from *r* = 0.0861 to *r* = 0.2553 at 85.61% coverage (seed 42, 32 unfiltered calibration genes), and 33-gene 10-fold spatial cross-validation reached *r* = 0.2845 and 81.03% coverage for *Acta2* (seed 444, 31 calibration genes after TISSUE’s default 1% *Flt1* filter, with overall 33-gene coverage of 79.05% at nominal 77% and 81.33% at nominal 80%, Supplementary Table S4).

### Pathogen and Vector Genomics Reanalyses on UseGalaxy.org

For the 30-sample *Aspergillus fumigatus* itraconazole time-course of Hokken et al. [26] (BioProject PRJNA482512, three clinical isolates sampled at 0, 30, 60, 120, and 240 minutes in duplicate), BRC-Analytics staged the Af293 reference genome (Ensembl Fungi ASM265v1) and 30 paired-end SRA runs into a UseGalaxy.org history (Supplementary Table S3). Workflow Foundry compiled a 60-step workflow chaining `fastp`, `rna_star` (`–quantMode` `GeneCounts`), unstranded count extraction (verified by forward-to-reverse STAR strand column ratios of 0.979–1.007 across ten libraries), and pooled 30-sample one-factor five-level timepoint `deseq2` (`∼timepoint`, HIDs 961 and 4423). Under the published study’s four simultaneous significance thresholds (baseMean ≥ 30, raw *P <* 0.05, *p*_adj_ *<* 0.1, and | log_2_ FC| *>* 1, where *P <* 0.05 is redundant with *p*_adj_ *<* 0.1 here and applying these thresholds directly to Hokken et al.’s Table S3 yields 3,968, 3,433, 1,205, and 1,095 DEGs vs. 3,955, 3,430, 1,207, and 1,101 reported in their table text), four separate 12-sample pairwise fits (HIDs 953– 959) inflated standard errors by a median of 20–26% and called 31% and 39% fewer differentially expressed genes at 120 and 240 minutes (missing 34% and 42% of the published DEG sets by overlap and recovering only 82 of 186 consistent DEGs). By contrast, the pooled 30-sample model matched the published standard error distribution (median ratio 1.000, interquartile range −0.4% to +0.3%, log_2_ FC Pearson *r* = 0.9996–0.9999). It identified 3,922 vs. 3,955 DEGs at 30 minutes, 3,393 vs. 3,430 at 60 minutes, 1,191 vs. 1,207 at 120 minutes, and 1,098 vs. 1,101 at 240 minutes, recovering 185 consistently differentially expressed genes against the published 186 (181 shared). Three custom User-Defined Tools (`hokken_go_enrichment`, `hokken_venn_comparison`, and `hokken_revigo_summary`) replaced the decommissioned AspGD GO Term Finder and automated cross-namespace `AFUA_/Afu` Venn overlap comparisons (benchmarked against BioVenn) and REVIGO semantic summarization inside Galaxy. Together, the three UDTs reproduced both the temporal enrichment inversion (135 upregulated vs. 2 downregulated terms at 30 minutes, reversing to 0 upregulated vs. 136 downregulated terms at 240 minutes) and the expression trajectories of the 61-gene drug-efflux transporter panel (*mdr1, mdr4, abcA, abcC*, and *mfs56*).

For the *Anopheles gambiae* infection chromatin study of Ruiz et al. [27] (GEO GSE152924, `SRR12064732–SRR12064735`), reads were aligned to AgamP4 with Bowtie2, filtered for MAPQ ≥ 10, and deduplicated. Sub-nucleosomal fragments (*<* 130 bp) were shifted by +4*/* − 5 bp via UDT `ruiz_tn5_fragment_extractor` and called with historical MACS2 2.1.2 via UDT `ruiz_macs2_212_callpeak` (`-f BAMPE -g 273109044 -q 0.01 -B –keep-dup all –nomodel –nolambda`, along-side single-end shifted `–shift -100 –extsize 200` sensitivity checks on the Galaxy MACS2 2.1.1 wrapper, Supplementary Table S3). Measured at the stated two-sided 51% overlap criterion (`bedtools intersect -f 0.51 -F 0.51`) against the 111,586 replicate-shared rows in the authors’ deposited table, independent MACS2 2.1.2 calls recovered 92.4% and 93.2% of reported midgut replicate-shared regions and 90.7% and 93.2% of reported salivary-gland replicate-shared regions (and 81.8–92.2% across all 193,005 deposited per-replicate peaks). Although Ruiz et al. refer to the 43,010 midgut and 68,576 salivary-gland replicate-shared rows as unique regions, the deposited GEO table lists each replicate’s peak as a separate unmerged row, so counting merged consensus intervals on strict 50–130 bp MACS2 2.1.1 peaks yielded only 9,448 midgut and 19,502 salivary-gland intervals (or 21,841 and 31,513 under relaxed calling and 21,775 and 30,844 without deduplication). Applying the stated two-sided 51% overlap rule row-wise per replicate to unmerged *<* 130 bp peaks via UDT `ruiz_reproducible_ths` recovered 41,580 midgut (−3.3%) and 57,518 salivary-gland (−16.1%) rows on MACS2 2.1.2 (and 41,340 [−3.9%] midgut and 57,274 [−16.5%] salivary-gland rows on the MACS2 2.1.1 wrapper), matching midguts closely while under-calling salivary glands. Fitting a single empirical near-containment threshold (`either_frac` = 0.851, where either peak covers at least 85.1% of the other, via UDT `ruiz_either_frac_shared_peaks` with downstream VectorBase rel49 gene, promoter, and enhancer annotations computed on the exported peak sets) reproduced 44,780 midgut (+4.1%) and 67,594 salivary-gland (−1.4%) rows along with assigned genes, promoters, and candidate enhancers derived from Ruiz Table S3 within 0.4–2.9% (0.4–4.6% across both tracks). A cross-tissue transfer test (`ruiz_either_frac_crossfit`) with pre-specified decision bands returned an explicit `TRANSFER INCONCLUSIVE` verdict (transfer error 7.16% vs. 5.0% threshold) because midguts and salivary glands exhibited distinct single-tissue optima (0.884 vs. 0.821).

For the *Aedes aegypti* midgut single-cell atlas of Wang et al. [28] (`SRR26591148` and `SRR26591149`, adult female midguts maintained on 10% sucrose), 10x Genomics 3’ v3 reads were aligned to AaegL5.0 (`GCF_002204515.2`, VectorBase-68 annotation) with `rna_starsolo` and filtered with `dropletutils` (`emptyDrops lower=500`). Whereas Wang et al. ran Cell Ranger v7.2.0 on VectorBase-65 (AaegL5.3), Seurat DoubletFinder, and Scanpy (*<* 30% mitochondrial reads using a hard-coded list of 38 AAEL mitochondrial gene identifiers), generic Scanpy quality-control tools in Galaxy match mitochondrial features by gene-symbol prefix. In our Galaxy history, prefix ND matched zero genes and terminated Scanpy quality control with `KeyError`: ‘`mito`‘, while an empty prefix flagged 100% of genes as mitochondrial. Prefixing the 38 mitochondrial genes with `MT_` in a custom GTF resolved the failure (100–2, 500 genes/cell and *<* 20% mitochondrial reads, without a separate doublet-removal step), retaining 6,707 post-QC cells (2,871 in replicate 1 `SRR26591149` at Louvain resolution 0.75 vs. 4,350 published, and 3,836 in replicate 2 `SRR26591148` at resolution 0.25 vs. 2,797 published, totaling 6,707 vs. 7,147 published). Cells were grouped into three conserved broad populations across 5,953 cells (`EC/digestive` `broad`: 3,331 and 2,004 cells, `HC-like/epithelial:` 73 and 213 cells, and `VM-like:` 257 and 75 cells, with 754 cells in minor replicate-specific clusters). Sample-blocked pseudobulk `edgeR` (∼0 + `coarse_label` + `sample_id` across *n* = 2 biological replicates, 6 pseudobulk profiles) across 8,248 expressed genes identified 1,784, 943, and 854 differentially expressed genes (FDR ≤ 0.05) across the pairwise contrasts. Pseudobulk testing recovered Table S1 markers including enterocyte marker `AAEL027580` (rank 1 in enterocyte vs. hemocyte-like, FDR = 2.2 × 10^−6^), visceral muscle markers `AAEL024504` (rank 1 in enterocyte vs. visceral muscle), `AAEL012704` (rank 1 in hemocyte-like vs. visceral muscle and rank 21 in enterocyte vs. visceral muscle), and muscle actin `AAEL001673` (rank 250, FDR = 0.002). It also recovered canonical midgut digestive proteases including zinc carboxypeptidase (`AAEL001863`, rank 309) and female-specific chymotrypsin (`AAEL003060`, rank 531, FDR *<* 0.001, Supplementary Table S3).

### Structure-Aware Phylogenomics of *Plasmodium vivax* VIR Antigens

Annotated proteomes, GFF3 gene models, and genome assemblies for *Plasmodium vivax* isolates PvP01 (Indonesia, short-read Illumina assembly with improved subtelomere scaffolding [30]), PvW1 (Thailand, long-read assembly [31]), and PvPAM (Peru, long-read assembly [32]) were screened against Pfam profile PF05795 using HMMER 3.4 (`hmmer_hmmsearch/3.4+galaxy0`) alongside six-frame pseudogene rescue via UDT `anton-1-gff_pseudogene_translator`. Domain screening identified 2,664 PF05795-positive VIR proteins (931 PvP01, 748 PvW1, and 985 PvPAM), of which 2,592 functional VIR proteins (897 PvP01, 735 PvW1, and 960 PvPAM) passed length filtering into structure-aware clustering. Sequence-only baseline clustering used MMseqs2 v18.8cc5c (`-e 1e-5 –min-seq-id` 0.3 -c 0.5) and MCL v22.282 (*I* = 2.0), producing 203 sequence clusters (146 subfamilies with ≥ 3 members, of which 132 subfamilies holding 94.8% of proteins already span all three genomes). Structure-aware 3Di embedding and clustering used Foldseek v10.941cd33 with ProstT5 weights (`-c 0.8 -e 0.001 –alignment-type 2`), producing 73 structural classes (24 with ≥ 3 members, dominant superfamily FS_001 = 1, 197 proteins, 46.2% of the repertoire, merging 89 sequence subfamilies) on the verified local CPU reference branch. On UseGalaxy.org, the same tool produced 76 structural classes across two independent runs on an ACCESS-CI NVIDIA A100 GPU via UDT `anton-gpu-xl-foldseek_prostt5_clustering` (25 classes with ≥ 3 members, FS_001 = 812 proteins, 336 sub-cluster representatives) and 76 structural classes on a 32-core Galaxy CPU node via UDT `anton-32-foldseek_prostt5_clustering` (30 classes with ≥ 3 members, FS_001 = 766 proteins). Because the two GPU runs matched identically while the local and Galaxy CPU runs differed (73 vs. 76 classes), the minor split reflects container environment or multi-threaded clustering tie-breaking rather than GPU versus CPU hardware. Across all branches, 20 to 21 structural classes contained members from all three continental genomes and held 96.5–97.3% of all VIR proteins (21 classes holding 2,521 of 2,592 proteins [97.3%] on local CPU, 20 classes holding 2,508 of 2,592 proteins [96.8%, or 97.6% of the 2,542 proteins in classes with ≥ 3 members] on Galaxy GPU, and 21 classes holding 2,501 of 2,592 proteins [96.5%] on Galaxy CPU), whereas 43 to 50 single-genome classes accounted for 2.0–2.7% of the repertoire (Supplementary Table S5). Lockstep amino acid and 3Di multiple sequence alignment of FS_001 sub-cluster representatives (*n* = 338 local: 118 Indonesia [34.9%], 110 Thailand [32.5%], and 110 Peru [32.5%], and *n* = 336 GPU branch, subsampled via MMseqs2 `–min-seq-id 0.5 -c 0.8 –cov-mode 0`) was performed with FoldMason v4.dd3c235 inside custom container quay.io/nekrut/foldmason-prostt5:4.dd3c235-cpu (UDT `anton-32-foldmason_align`). Column-occupancy trimming (trimAl -gt 0.65) on the Galaxy branch retained 176 of 336 representatives whereas the verified local CPU reference branch retained all 338 representatives. Partitioned maximum-likelihood phylogenetic inference in IQ-TREE 3 (amino acid VT+F+R6 and 3Di Garg-Hochberg models, 1,000 ultrafast bootstrap replicates) and 1,000-shuffle Fitch parsimony permutation testing (observed score = 160 vs. null median 160, mean 159.2 ± 3.8, range 145–177, *p* = 0.55) are therefore reported from the 338-tip local CPU reference branch. Among the 1,672 VIR genes placed on the 14 main chromosomes (of 2,664 total PF05795-positive genes, with the remaining 992 located on unplaced subtelomeric scaffolds), 1,098 (65.7%, or 64.5% of chromosome-placed functional VIR genes, ranging from 57.1% in PvW1 and 58.3% in PvPAM to 89.5% in PvP01) fell within the outer 10% of chromosome arms (`anton-1-spatial_analysis`). Flanking microsynteny analysis (`anton-8-mmseqs_easy_rbh_pair and anton-1-microsynteny_filter`) showed that 11 of 194 local (3 of 141 GPU) FS_001 reciprocal best-hit pairs retained synteny, compared against 5,120 single-copy housekeeping reciprocal best hits (median amino acid identity 99.8%, Supplementary Table S5).

### Manifold Emergence Dating Across 42 Pathogen Cohorts

We executed the 42-cohort molecular clock study in Orbit across two separate prompt-driven stages on UseGalaxy.org (Table 3, full prompt files archived in `manuscript/prompts/`). In the first stage, a dedicated harvesting and preparation prompt (`prompt-galaxy.md`, adapted from our autonomous BEAST XML harvester prompt `prompt.md`) directed Orbit to inventory, fetch, and validate 42 self-contained author BEAST v1.x and v2.x XML configurations from GitHub, Zenodo, Dryad, and Figshare across 37 primary publications (41 distinct pathogen outbreaks plus a paired 14,517 bp coding tutorial partition, `41_skygrid_rabies_gill2020`, for the 2014 Sierra Leone Ebola cohort [36, 48]), preserving original binary bytes (`chronaeon_reference_42_originals.tar.gz`) and parsing inlined nucleotide alignments, verifiable tip dates, clock specifications, and linked publication DOIs via UDT `chronaeon_batch_preparator` v1.0.0 into synchronized 42-element BEAST XML, aligned FASTA, and tip-date collections in preparation history `bbd44e69cb890` `6b5c7c1fd5515b95301` (Supplementary Table S6). In the second stage, a companion analysis prompt (`prompt-galaxy-analysis.md`) directed Orbit to execute concurrent manifold emergence dating across all 42 prepared collections in analysis history `bbd44e69cb8906b55342b6485eac7` `7f7` using UDT `chronaeon_cohort_analysis` v3.2.2 [38], completing in seconds of core manifold computation per alignment (for example, 2.58 seconds for TN93 AutoClock manifold regression in pilot benchmarking, compared with 5 × 10^6^ to 10^9^ MCMC states taking 8–36 hours in dual-core BEAST X or 5–7 days in BEAST 2.7.7). During the initial 42-element batch execution on UseGalaxy.org, 5 of 42 container jobs hit a 1-minute sed timeout, were re-executed across recovery runs, and were merged into the final 42-element output collections via `BUILD_LIST` . For each alignment, the tool computed pairwise evolutionary distances, continuous manifold embeddings, analytical Fieller 95% confidence intervals [49], leave-one-out cross-validation error, AutoClock normalized Laplacian spectral bisection (*K*^*^), and optional HyphAeon Attention PGLS root inference using offline-staged `model.viral.pt` weights.

**Table 3:**
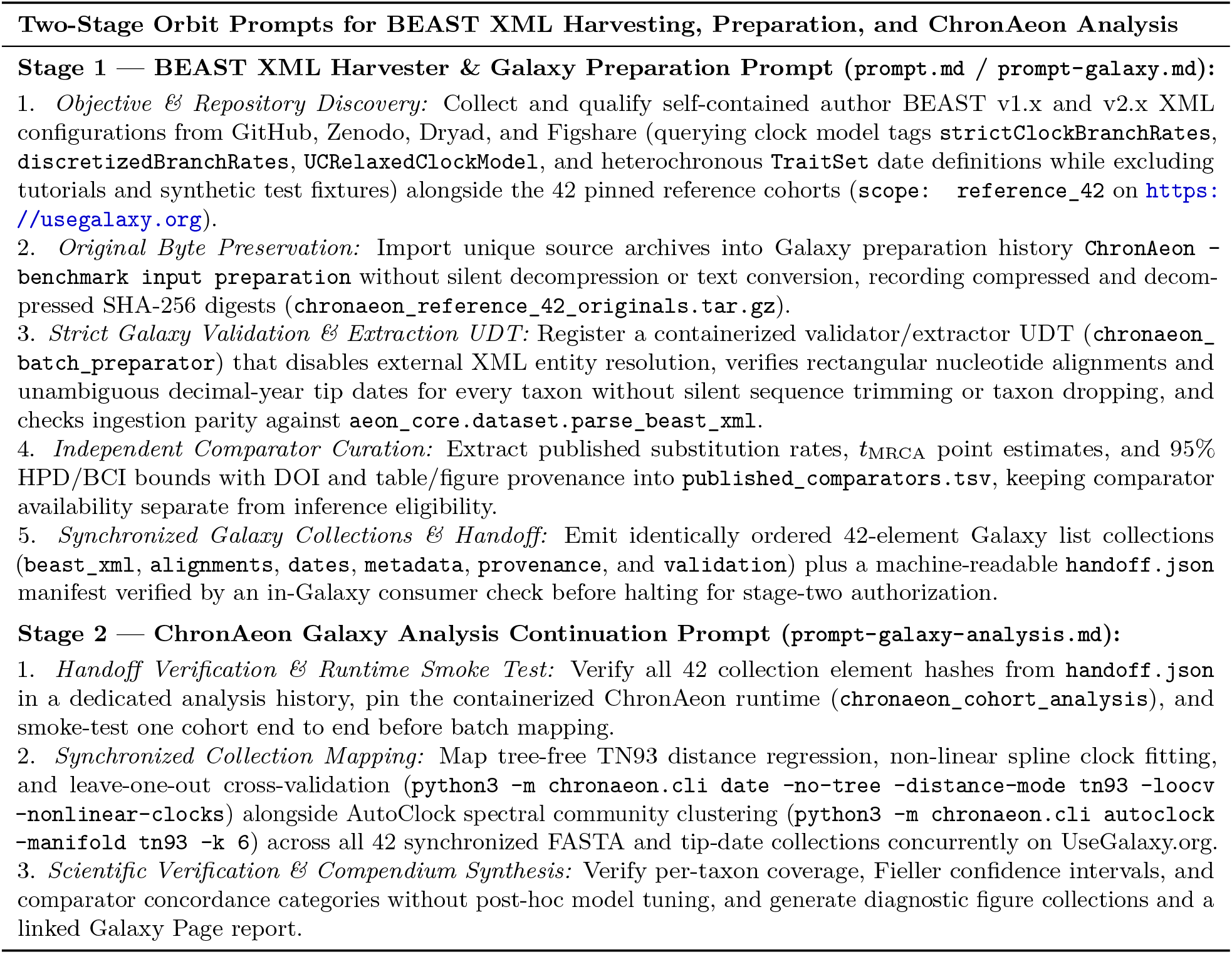
Summary of the two-stage Orbit prompts used to harvest, validate, and analyze the 42 BEAST XML pathogen cohorts on UseGalaxy.org. Full verbatim prompt files (`prompt.md`, `prompt-galaxy.md`, and `prompt-galaxy-analysis.md`) are archived in `manuscript/prompts/`.

To audit reference comparator provenance across all 42 alignments (`chronaeon_42cohorts_ provenance.tsv` and Supplementary Table S7), we classified each cohort’s reference interval into three categories. First, 15 alignments had explicit 95% highest posterior density (HPD) or Bayesian credible intervals (BCI) reported in the primary publication’s text or tables (`plus 39_chikv_ reunion_dellicour2020` [50], which carries a published BEAST root of 2004.8 [2004.5, 2005.1] in published_comparators.tsv but was marked `NO_COMPARATOR` in the automated runner). Second, 24 alignments had numeric reference intervals in `aggregate_benchmark_results.tsv` reconstructed from prose approximations (such as Hill and Baele’s “approximately March 2014” TempEst intercept [36] bracketed as 2014.20 [2014.10, 2014.30], or O’Toole et al.’s late-2015 APOBEC3 emergence estimate [51]), molecular clock calibrations, or Skygrid grid cutoffs. Third, 2 additional alignments were unscored (`NO_COMPARATOR`) in the automated runner and carried prose clade ranges in `published_comparators.tsv: 38_mab_commins2023` (*Mycobacterium abscessus* [52], prose range ∼ 1960–1980 vs. ChronAeon 1975.87 [1966.21, 1982.46]) and `40_hiv1_crf01ae_philippines2024` (HIV-1 CRF01_AE Philippines, prose range ∼ 1995–2002 vs. ChronAeon 1975.84 [1969.29, 1980.93], alongside 39_chikv_reunion_dellicour2020 where ChronAeon inferred 1954.99 [1948.09, 1960.62]). In addition, cohort 41_skygrid_rabies_gill2020 (14,517 bp Ebola coding partition) duplicated the summary row of cohort `00_ebola_sierraleone_gire2014` (4,453 bp variable-site partition) in `aggregate_benchmark_results.tsv`, whereas the single-pass rerun in `deterministic_benchmark_ results.tsv` recorded its distinct 14,517 bp estimates (2012.71 OLS and 2002.60 spline). We therefore report benchmark concordance and median errors both across all 42 rows (39 numeric comparators) and across the 41 unique alignments (38 unique numeric comparators).

On the 196-taxon 2014 Sierra Leone Ebola benchmark (cohorts 00 and 41 [34, 36]), the reconstructed tutorial origin is 2014.20 [2014.10, 2014.30] and the Gire et al. 81-sequence published BEAST root is 2014.14 [2014.07, 2014.20]. Galaxy TN93 OLS estimated a root of 2014.23 (95% Fieller CI: [2014.17, 2014.27], cross-validation MAE: 26.4 days, portal panel: 2014.30 [2014.24, 2014.34], cubic spline: 2004.92). For the 1,610-taxon West African Ebola Makona epidemic [35], the multi-model suite inferred 2013.81 ([2013.74, 2013.86], portal panel: 2013.57 [2013.47, 2013.66]), overlapping the published BEAST 95% HPD of 2013.98 ([2013.79, 2014.15]). For SARS-CoV-2 Gamma (P.1) [37], TN93 OLS inferred 2020.71 (95% Fieller CI: [2020.63, 2020.76], MAE: 17.0 days, portal PGLS panel: 2020.61 [1595.27, 2020.79]) against the published BEAST root of 2020.87 ([2020.78, 2020.93]). Because the OLS Fieller interval lies just below the published 95% HPD without overlapping, it qualifies as concordant by point-estimate proximity (|Δ*t*_MRCA_| = 0.16 *<* 1.0 year). On the single-pass portal baseline plotted in Figure 3, 18 of 39 numeric-comparator cohorts (46.2%, or 42.9% of 42) were visually concordant (interval overlap or |Δ*t*_MRCA_| *<* 1.0 year) and 21 (50.0% of 42) diverged. Under best-of-model selection across pure TN93 OLS and cubic spline regression in `aggregate_benchmark_results.tsv`, ChronAeon achieved concordance across 24 of 42 alignments (57.1%, or 61.5% of 39 numeric comparators, and 23 of 41 [56.1%] or 23 of 38 [60.5%] unique cohorts). Switching between OLS and spline in `deterministic_ benchmark_results.tsv` shifts estimates across 29 of 42 cohorts. The 24 concordant rows comprise 13 alignments (12 unique, including Zika Angola 08 at |Δ*t*_MRCA_| = 0.79 years: 2005.71 [2000.71, 2007.45] vs. 2006.50 [2005.80, 2007.20]) with |Δ*t*_MRCA_| *<* 1.0 year and 11 cohorts qualifying by interval overlap alone. Interval-overlapping cohorts included Caribbean DENV-1 04 (1846.46 [1830.56, 1859.76] compared with BEAST 1880.98 [1844.11, 1901.82]) [53], DENV polyprotein 06 (1969.33 [1966.02, 1972.11] vs. 1965.00 [1958.00, 1972.00]), and DENV-1 07 (1940.17 [1892.32, 1956.84] vs. 1952.00 [1945.00, 1960.00]). Additional interval-overlapping cohorts were Influenza H3N2 12 (1966.63 [1965.56, 1967.63] vs. 1968.00 [1967.50, 1968.50]) [54], Influenza H5N1 13 (1993.14 [1992.09, 1993.84] vs. 1994.50 [1993.80, 1995.80]), Caribbean DENV-2 05 (1842.02 [1592.75, 1957.09]), and *Y. pestis* second pandemic 29 (1141.78 [−383.82, 1335.47]) [55]. Restricted to the 15 cohorts with explicit published 95% HPDs/BCIs in the primary text, concordance was 10 of 15 (66.7%, or 10 of 16 [62.5%] including `39_chikv_reunion_dellicour2020`). Pairing pairwise-distance regression with HyphAeon Attention PGLS (model.viral.pt) resolved structured cohort `30_mpox_clade_ib_burundi2025` (2024.29 [2024.19, 2024.36] vs. reference 2023.95 [2023.67, 2024.22], |Δ*t*_MRCA_| = 0.34 years, where unguided spline extrapolation gave 1967.29). Attention PGLS raised multi-model concordance to 25 of 42 alignments (59.5%, or 64.1% of 39, and 24 of 41 [58.5%] or 24 of 38 [63.2%] unique cohorts). Across all 39 numeric-comparator rows, median |Δ*t*_MRCA_| was 7.19 years for pure pairwise-distance regression and 6.41 years with Mpox PGLS (8.32 and 6.80 years across the 38 unique comparators). Across the 19 post-1980 comparator rows, median |Δ*t*_MRCA_| was 0.51 years for pure pairwise-distance regression and 0.40 years with Mpox PGLS (0.65 and 0.46 years across the 18 unique post-1980 cohorts). Among the 14 discordant comparator cohorts, several reflected ancient roots compressed by purifying selection and saturation, such as Lassa (`18_lassa_andersen_suchard2020`) [43,44], *Mycobacterium tuberculosis* (35–36) [56,57], and Measles/Rinderpest (`37_measles_1912_dux2020`, unbounded Fieller interval [−∞, 1870.69] [49] and portal interval [944.09, 1380.77] vs. 528 BCE [−528 CE] root [45]). The remainder were multi-lineage alignments where whole-alignment regression dated a deep species split instead of a focal regional introduction, including Zika Pacific/Fiji (14), Avian Influenza H7/N7 (19–20), Caribbean DENV-3/4 (23–24), Foot-and-Mouth Disease serotype A (25), Mpox Clade I DRC (27) [58], RSV-A (31), and African Swine Fever (34) (Supplementary Tables S6 and S7).

## Data and Code Availability

Primary sequencing datasets and genome assemblies analyzed in this study are available from public repositories. Bacterial RNA-seq runs (`ERR14660412`, `ERR14660413`, and `ERR14660420– ERR14660422`) are under ENA BioProject `PRJEB86181`, and the *B. cereus* ATCC 10987 reference assembly (`GCF_000008005.1` / `GCA_000008005.1`) is under NCBI BioProject PRJNA74. Fungal, vector chromatin, and vector single-cell datasets are available under NCBI BioProject PRJNA482512 (*A. fumigatus* Af293 `ASM265v1`), NCBI GEO accession GSE152924 (`SRR12064732–SRR12064735`), and NCBI SRA runs SRR26591148–SRR26591149 (*A. aegypti* `GCF_002204515.2`). Spatial and phylogenetic benchmark archives are available at Zenodo record `8259942` (TISSUE Dataset 15) and on UseGalaxy.org (42 BEAST XML benchmarks from https://github.com/veg/chronaeon and https://github.com/veg/HyphAeon, Supplementary Tables S6 and S7). Galaxy history, workflow, collection, and Page links for each investigation are provided in Supplementary Tables S3–S6.

All core platform software components developed in this work are open-source and publicly available:

- Galaxy MCP Server: https://github.com/galaxyproject/galaxy-mcp.
- Orbit Workbench and Loom Runtime: https://github.com/galaxyproject/loom.
- Galaxy Workflow Foundry: https://github.com/galaxyproject/foundry.
- BRC-Analytics Portal: https://brc-analytics.org (repository: https://github.com/galaxyproject/brc-analytics).
- Galactic Skills and Developer Guides: https://github.com/galaxyproject/galaxy-skills.
- Universal Agentic Plugins: https://github.com/galaxyproject/agentic-plugins.

Manuscript figure scripts, in-repository figure source tables (`manuscript/figures/data/`), benchmark summary tables, and companion case-study repositories (https://github.com/nekrut/orbit-paper, https://github.com/nekrut/spatTrOrbit, https://github.com/nekrut/phist, https://github.com/veg/chronaeon, and the BRC reanalysis repositories in Supplementary Table S3) are archived on GitHub and UseGalaxy.org (with any histories, workflows, or companion repositories under review access controls made publicly accessible upon publication).

## Acknowledgments

We thank Anup Kumar, Ivo Leist, Wolfgang Maier, Daniela Schneider, and Paul Zierep of the University of Freiburg (UseGalaxy.eu) for helpful discussions, technical contributions, and feedback on the manuscript, and David Y. Hancock and the Jetstream2 team at Indiana University for supporting the cloud infrastructure where User-Defined Tools execute. This work was supported by National Institutes of Health grants U24AI183870 (NIAID) and U24HG006620 (NHGRI), and by National Science Foundation awards 2419522 and 2138585. This work used the Jetstream2 cloud resource at Indiana University [59] (supported by U.S. National Science Foundation award OAC-2005506), Stampede3, Lonestar6, and Ranch at the Texas Advanced Computing Center (TACC), Bridges-2 at the Pittsburgh Supercomputing Center, Expanse at the San Diego Supercomputer Center, and Rockfish at Johns Hopkins University through allocation MCB140147 from the Advanced Cyberinfrastructure Coordination Ecosystem: Services & Support (ACCESS) program [60], which is supported by U.S. National Science Foundation grants #2138259, #2138286, #2138307, #2137603, and #2138296. The authors acknowledge Indiana University and the Jetstream2 team (https://jetstream-cloud.org) and the Texas Advanced Computing Center (TACC, http://www.tacc.utexas.edu) at The University of Texas at Austin for providing high-performance computing, cloud, and storage resources that have contributed to the research results reported within this paper, alongside computational resources provided by UseGalaxy.org and UseGalaxy.eu.

## Supplementary Information

**Figure S1:**
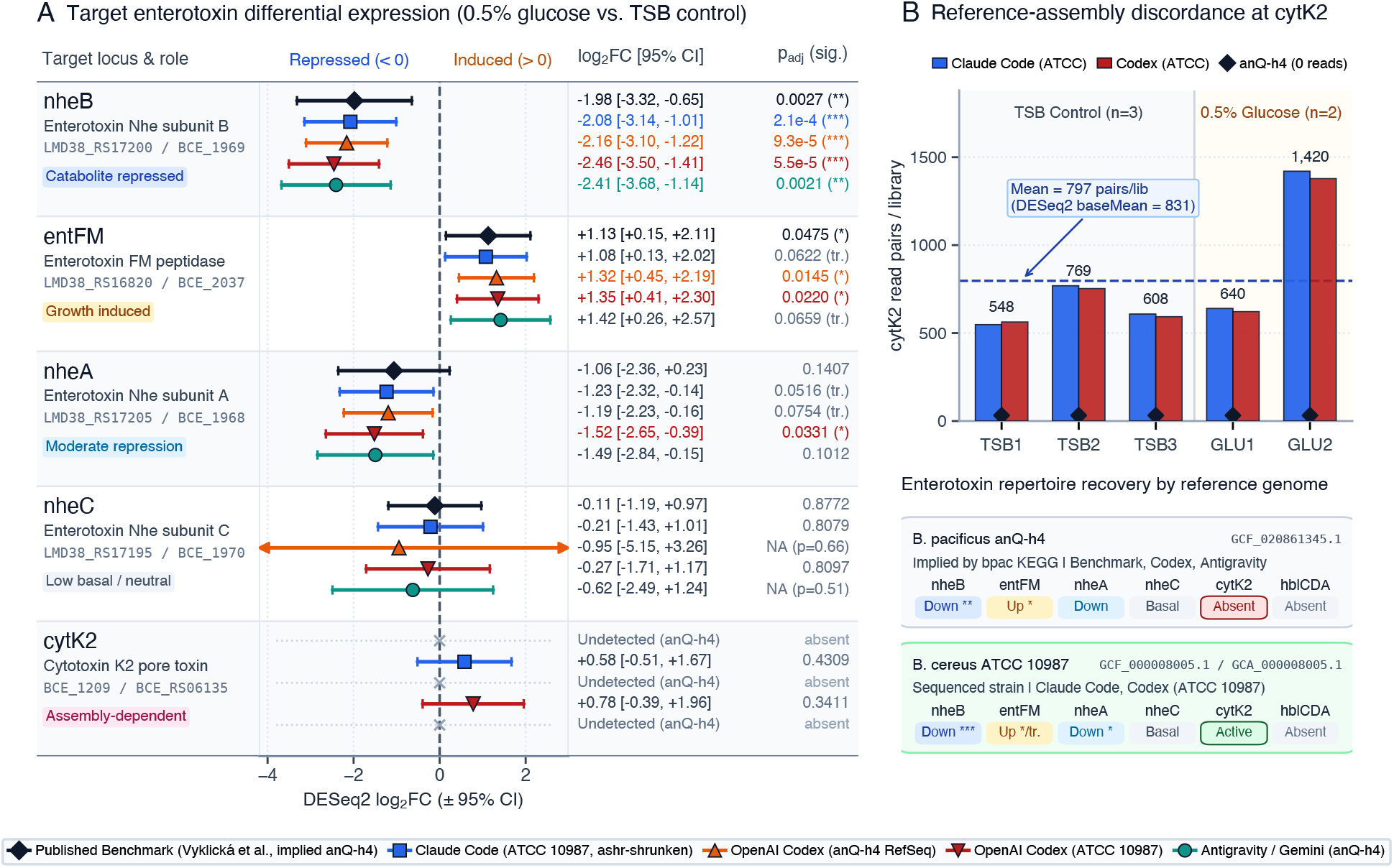
Local agent runs recover headline enterotoxin signals but diverge on reference genomes and leave unversioned glue code. **A**, Differential expression estimates (log_2_ FC with 95% confidence intervals) and adjusted *p*-values for target enterotoxin genes (*nheB, entFM, nheA, nheC*, and *cytK2*) compared against the published study (black diamonds, Vyklická et al. [15]). OpenAI Codex (orange and rust across two reference assemblies) and Google Gemini via Antigravity (teal) report unshrunken DESeq2 Wald estimates, whereas Claude Code (blue, `nf-core/differentialabundance`) reports `ashr`-shrunken estimates (for *nheC* on *B. pacificus*, where independent filtering sets *p*_adj_ to NA, the unadjusted *p*-value is shown). All three agents confirmed strong repression of *nheB* and induction of *entFM*. **B**, Reference-assembly mismatch at the cytotoxin K (*cytK2*) locus. Aligning against the sequenced isolate *B. cereus* ATCC 10987 in Claude Code and Codex recovered active baseline *cytK2* transcription across all libraries (548–1,420 read pairs per library, mean 797, DESeq2 baseMean = 831, *p*_adj_ = 0.43). Aligning against *B. pacificus* anQ-h4 missed *cytK2* entirely because that assembly lacks the locus.

**Figure S2:**
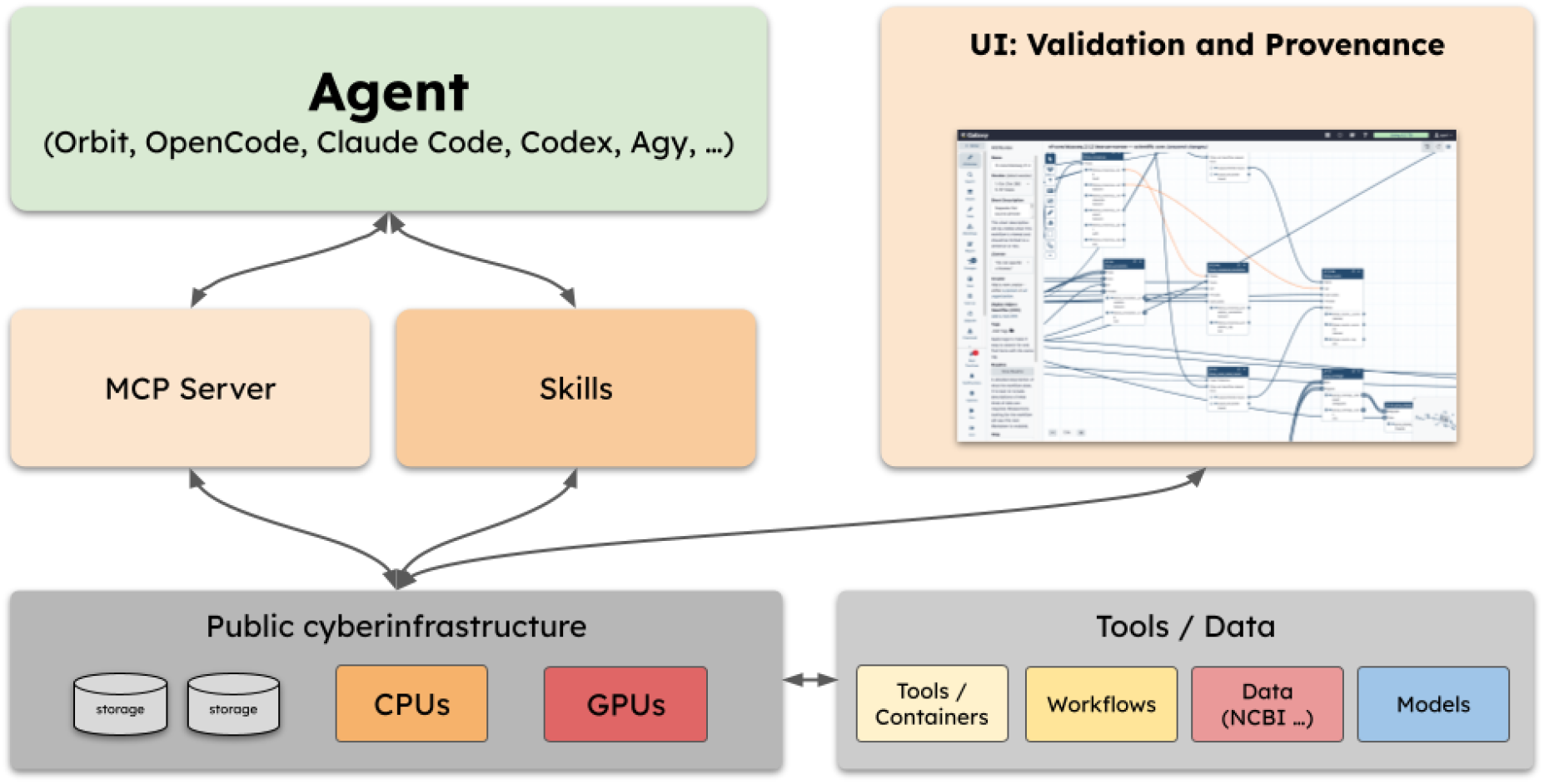
Architecture for verifiable agentic science on public infrastructure. **Agent** (top left): Researchers use Orbit or connect any generic coding agent (such as OpenCode, Claude Code, OpenAI Codex, or Antigravity CLI). **MCP Server and Skills** (middle left): Curated domain skills supply validated bioinformatics protocols, while the Galaxy MCP server translates agent actions into typed API calls in place of unconstrained shell commands. **UI: Validation and Provenance** (top right): Galaxy’s visual workflow editor and history interface provide pre-execution schema validation and an inspectable provenance record for human steering and verification. **Public cyberinfrastructure and Tools/Data** (bottom): Galaxy schedules containerized tools, peer-reviewed workflows, public datasets (NCBI, ENA, BRC-Analytics), and large models across CPU, GPU, and storage allocations on national supercomputing infrastructure.

**Figure S3:**
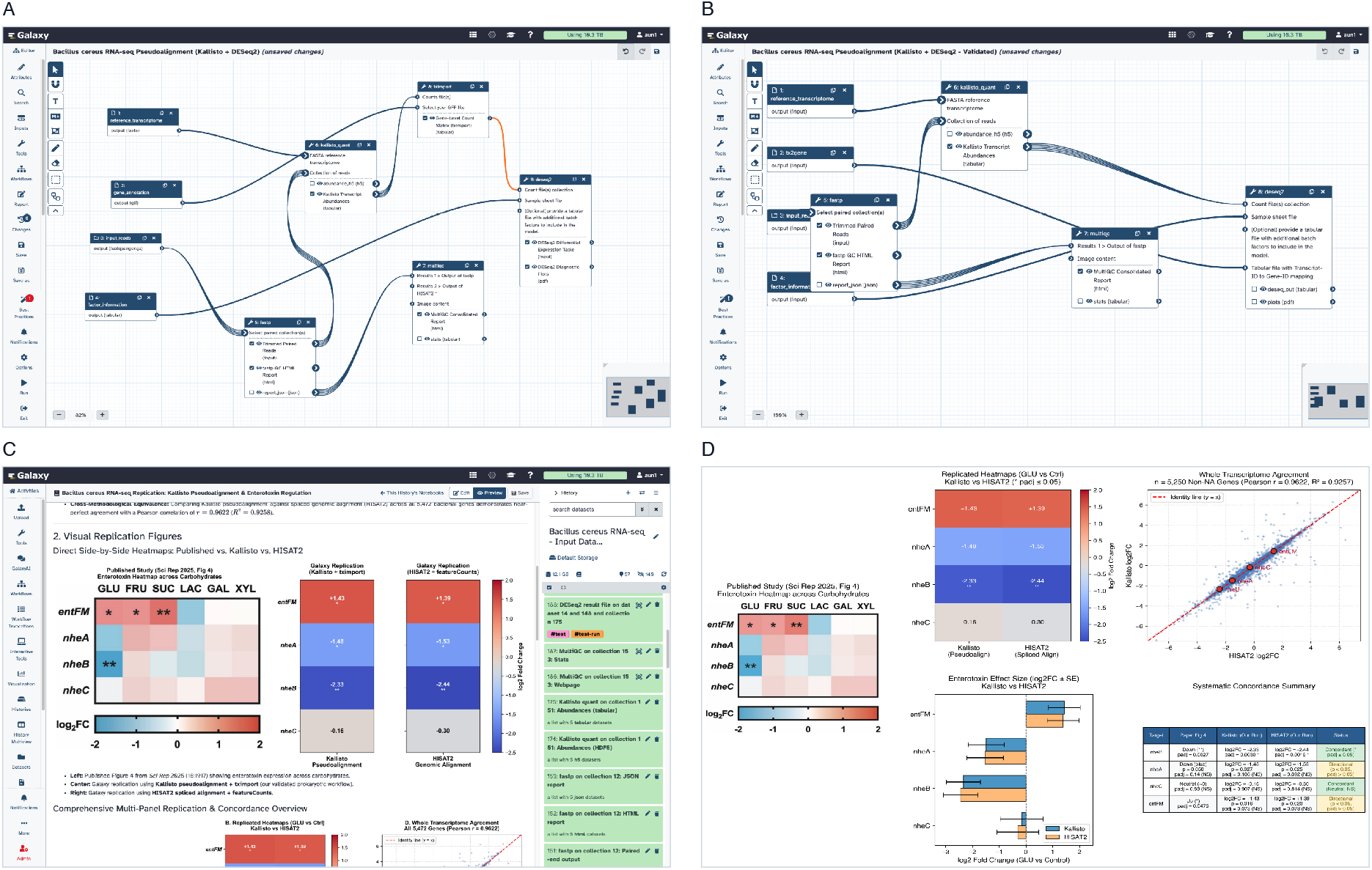
Workflow and execution provenance on free public infrastructure catches pipeline errors and reproduces bacterial RNA-seq profiles. **A**, Pre-execution workflow validation on UseGalaxy.org. When an agent connected via the Galaxy MCP server inserts an incompatible standalone `tximport` step between a Kallisto output collection and DESeq2, the workflow editor flags the type mismatch in orange. **B**, Validated workflow after repair. Passing the transcript-to-gene table to DESeq2’s built-in `tximport` mode resolves all workflow connections in green. **C**, Live provenance and interactive Galaxy History Notebook. Jobs execute on free public ACCESS-CI supercomputing allocations while completed datasets populate the right-hand history panel and the center notebook links narrative text to verified history dataset IDs. **D**, Biological replication of Vyklická et al. [15] across the *B. cereus* ATCC 10987 coding transcriptome. Supplementation with 0.5% glucose represses enterotoxin subunits *nheB* (5.05-fold down, log_2_ FC = −2.335, *p*_adj_ = 2.99 × 10^*−*3^) and *nheA* (2.80-fold down, log_2_ FC = −1.484, raw *p* = 0.0274, *p*_adj_ = 0.106), leaves component *nheC* unchanged (log_2_ FC = −0.158, *p*_adj_ = 0.907), and induces *entFM* (2.70-fold up, log_2_ FC = +1.432, raw *p* = 0.0158, *p*_adj_ = 0.0730). Across all 5,250 expressed coding loci (of 5,472 total), Kallisto pseudoalignment and HISAT2 alignment agree closely (Pearson *r* = 0.9622, *R*^2^ = 0.9257).

**Figure S4:**
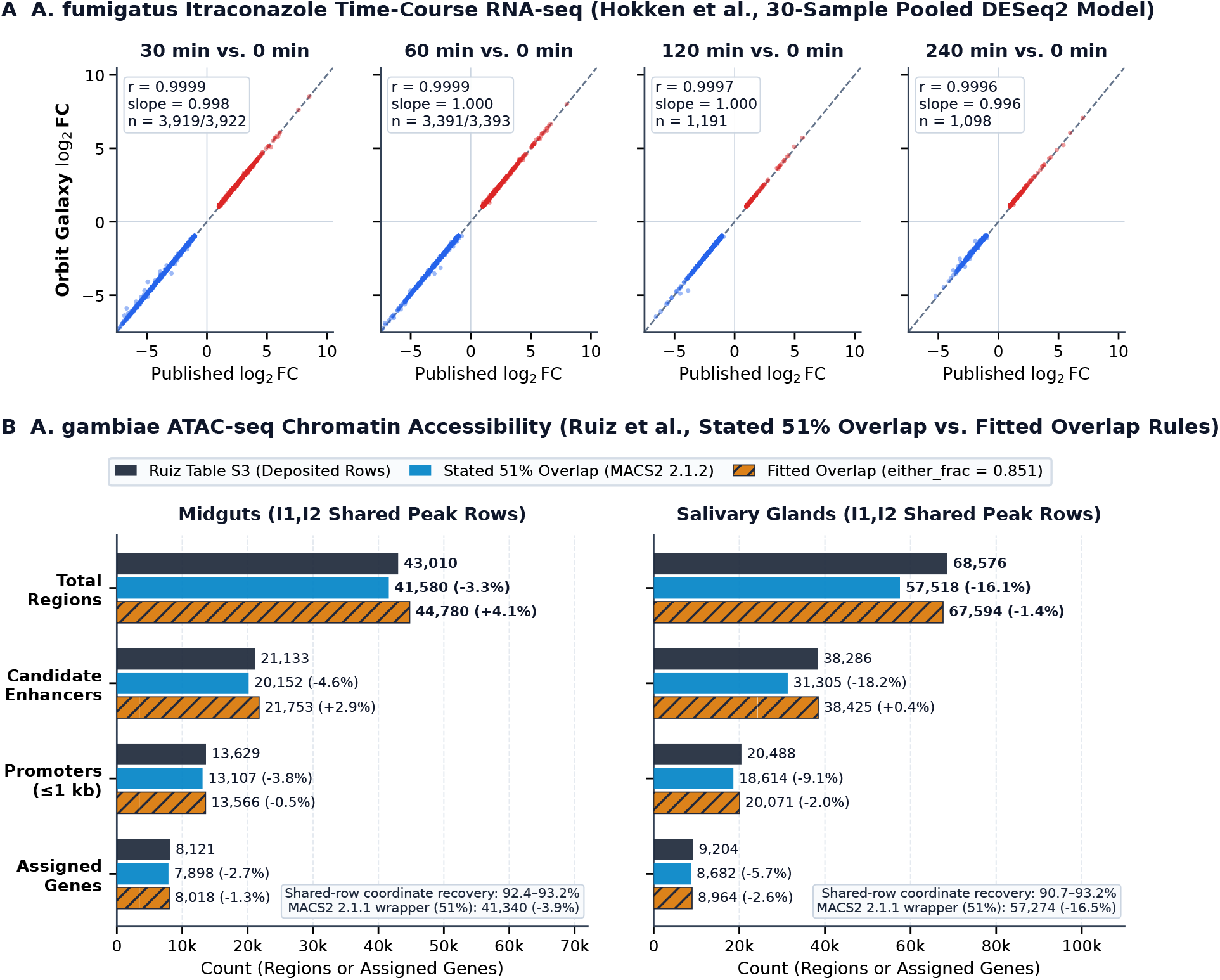
Reproducing pathogen and vector functional genomics studies on UseGalaxy.org. **A**, Transcriptomic response of the pathogenic mold *Aspergillus fumigatus* to the antifungal drug itraconazole across 30 libraries (Hokken et al. [26]). A pooled 30-sample DESeq2 model compiled via Workflow Foundry reproduces published log_2_ fold changes across all four post-exposure timepoints (*r* = 0.9996–0.9999, slopes 0.996–1.000). Published differentially expressed gene counts match within 0.3–1.3% (*n* = 3, 919 of 3, 922 at 30 min, 3, 391 of 3, 393 at 60 min, 1, 191 at 120 min, and 1, 098 at 240 min). **B**, Chromatin accessibility in *Anopheles gambiae* malaria mosquito midguts and salivary glands during *Plasmodium falciparum* infection (Ruiz et al. [27]). Independent MACS2 peak calling recovers 90.7–93.2% of the 111,586 replicate-shared genomic regions in the authors’ deposited table across all four libraries (81.8–92.2% across all 193,005 deposited peaks). Provenance-tracked overlap rules show how unmerged per-replicate rows and overlap filtering account for the headline region, promoter, and enhancer counts derived from Ruiz Table S3 within 0.4–2.9% (0.4–4.6% across tracks).

**Figure S5:**
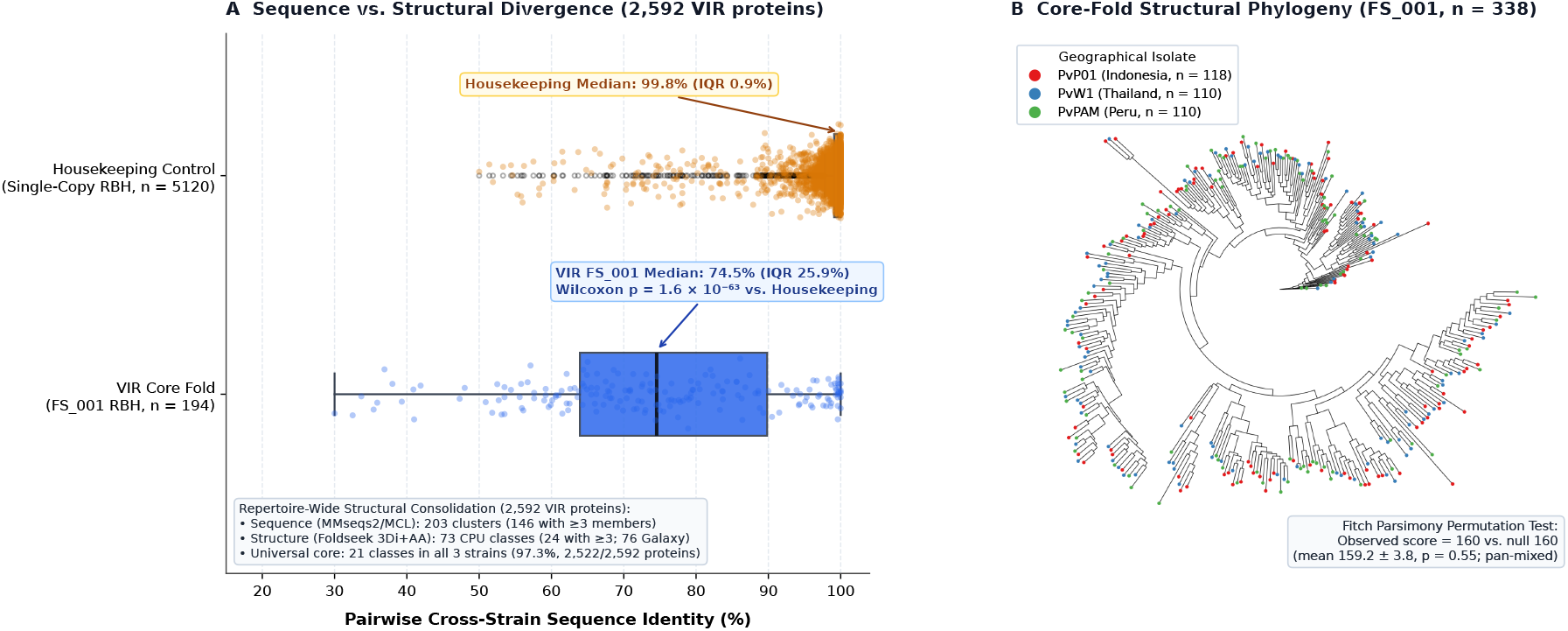
Structure-aware family reconstruction and lineage sharing of *Plasmodium vivax* VIR surface antigens across continental genomes. **A**, Sequence twilight zone and structural unification of over-split subfamilies. Horizontal box-and-strip plots display pairwise amino acid sequence identity between PvW1 (Thailand) and PvPAM (Peru) for conserved single-copy housekeeping genes (median 99.8%, IQR 0.9%, amber, *n* = 5, 120 pairs) versus VIR pairs within dominant superfamily FS_001 (median 74.5%, IQR 25.9%, Wilcoxon *p* = 1.6 × 10^*−*63^, blue, *n* = 194 pairs). Sequence-only Markov clustering at 30% minimum sequence identity splits the 2,592 VIR proteins across Indonesia (PvP01), Thailand (PvW1), and Peru (PvPAM) into 203 sequence clusters (146 subfamilies with ≥ 3 members). Structure-aware clustering using ProstT5 and Foldseek 3Di representations collapses this repertoire into 73–76 structural classes (24–30 with ≥ 3 members) across local and UseGalaxy.org execution environments, where the dominant superfamily FS_001 unifies 89 sequence subfamilies on the local reference branch (see Methods). **B**, Circular maximum-likelihood structural phylogeny inferred by IQ-TREE 3 from FoldMason alignments of 338 FS_001 representatives on the local reference branch across Indonesia (PvP01, red, 34.9%), Thailand (PvW1, blue, 32.5%), and Peru (PvPAM, green, 32.5%) using partitioned amino acid and 3Di structural models. Continental lineages are interleaved across the tree, and a tip-label parsimony permutation test shows no detectable geographic clustering (observed score = 160 vs. 1,000 random shuffles with null median 160, *p* = 0.55).

**Table S1:** Resource allocation, software environments, and compute footprints across three autonomous agent execution runs. All three runs re-analyzed the primary glucose versus control contrast from the *Bacillus cereus* transcriptomic benchmark (BioProject PRJEB86181) on an identical host machine (Ubuntu Linux workstation with 24 CPU cores, 62 GB RAM, and NVMe SSD storage). While all three platforms recovered the core biological signal of enterotoxin regulation, their resource management paradigms differed sharply. Gemini utilized a lightweight, greedy Unix shell pipeline with immediate file cleanup. Claude executed a heavy containerized Nextflow workflow that consumed over 100 GB of scratch disk and caused disk exhaustion until secondary quality control tools were disabled. Codex applied strict computational budgeting. It capped memory buffers and thread allocations per stream while maintaining an explicit ledger of purged intermediate files. None of the three runs produced persistent provenance records that external researchers could inspect or re-execute without rebuilding the local execution environment.

| Metric | Google Gemini 3.8 Flash (Antigravity) | Anthropic Claude Opus 5.5 (Claude Code) | OpenAI Codex / GPT-6 Astra (Codex CLI) |
| --- | --- | --- | --- |
| **Agent Runtime** | Antigravity CLI (`agy`) driving ad-hoc Bash and R scripts | Claude Code (`cc`) driving Nextflow workflows via Docker | Codex CLI (`codex exec`) driving modular Bash runner |
| **Software Stack** | Bowtie2 v2.5.5, feature-Counts v2.1.1, DESeq2 v1.50.2, fastp v1.4.0 | nf-core/rnaseq 3.14.0 (STAR, Salmon), nf-core/differentialabundance 1.4.0 | HISAT2 v2.2.1, feature-Counts, DESeq2, fastp v0.24.0, FastQC |
| **Thread & CPU Strategy** | Greedy 16-thread allocation across download, alignment, and quantification | Automated Docker scheduling across 24 host CPU cores | Strict budget: `OMP/MKL=1`, explicit slices (10 align, 3 sort, 2 QC, 4 count) |
| **Peak Memory (RAM)** | 2–4 GB (standard heap allocation) | 16–30 GB (STAR 2-pass indexing and alignment in Docker) | <6 GB (explicit buffer limit: `samtools sort -m 512M`) |
| **Disk Footprint** | 15–20 GB (immediate inline cleanup of raw SRA and FASTQ files) | >100 GB scratch disk (49 GB Nextflow cache and >40 GB QC sorting files) | <20 GB (sequential processing, pruned FASTQs tracked in ledger) |
| **Dependency Model** | Shared host Conda environment (`bacillus_rnaseq`) | Local Conda environment plus Docker OCI container layers | Project-isolated cache (`.cache/`) with explicit package URLs |
| **Custom Scripts Generated** | 3 scripts (1 Bash, 1 Python, 1 R, 302 LOC) | 9 scripts (7 Bash, 1 Python, 1 R, 190 LOC) | 14 scripts (6 Bash, 7 Python, 1 R, 774 LOC) |
| **Reference Strategy** | Single reference (*B. pacificus* anQ-h4 RefSeq) | Single reference (*B. cereus* ATCC 10987 GenBank/RefSeq) | Dual-reference sensitivity (*B. pacificus* anQ-h4 and ATCC 10987) |
| **Execution Friction** | None (completed without intervention) | Host disk exhaustion (required manual container deletion and skipping QC) | None (preventative buffer limits and resumption caching) |

**Table S2:**
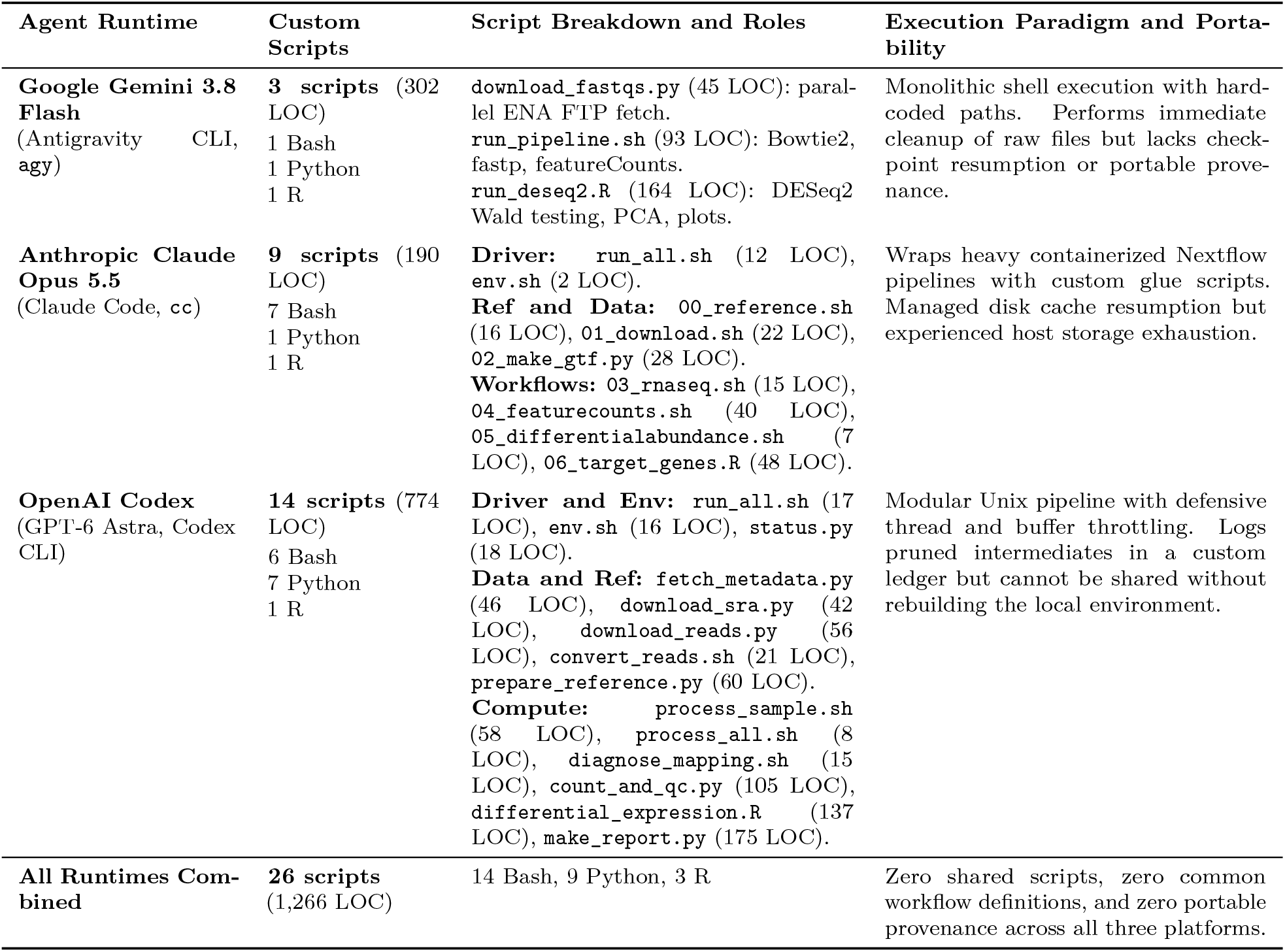
Summary of custom scripts generated by autonomous agents during empirical reanalysis. Number of custom scripts, programming languages, code volumes, and functional scopes generated by each runtime to replicate the *Bacillus cereus* RNA-seq benchmark. Gemini implemented a minimal three-script pipeline driving standard command-line utilities. Claude built nine wrapper and glue scripts to drive containerized nf-core workflows and parse downstream abundances. Codex constructed a modular fourteen-script suite featuring automated metadata queries, dual-reference mapping, explicit buffer throttling, and HTML dashboard synthesis. None of the twenty-six scripts were portable or auditable beyond the specific workstation where they were generated.

**Table S3:** Computational artifacts from the bacterial, fungal, and vector genomics reanalysis studies. Galaxy histories, workflows, User-Defined Tool definitions, and interactive Galaxy Pages archived on UseGalaxy.org.

| Study / System | Identifier / Link | Description |
| --- | --- | --- |
| **Vyklická et al. (2025)**<br>*Bacillus cereus* RNA-seq<br>(PRJEB86181) | bbd44e69cb8906b59549f5524d3906f3<br>bbd44e69cb8906b5d979c5818e07f6ad<br>ea7670a8c3fa3562<br>93fcddc692e0986f | Complete 5-library prokaryotic RNA-seq replication history on *B. cereus* ATCC 10987 (raw CDS HID 66, deduplicated CDS HID 106, `tx2gene.tsv` HID 148, Kallisto collection HID 175, DESeq2 results HID 188, and JupyterLab comparison notebook HID 192), initial HISAT2/featureCounts comparison history (HIDs 1–64), validated four-step Kallisto–DESeq2 integrated `tximport` workflow, and published interactive Galaxy Page. |
| **Hokken et al. (2019)**<br>*Aspergillus fumigatus* RNA-seq<br>(PRJNA482512) | bbd44e69cb8906b5a5cd735af575e8f5<br>b3638784a5d2aa43<br>b4346d545fb2b437 | Complete 30-sample time-course history (pairwise DESeq2 fits HIDs 953–959, pooled 30-sample `~timepoint` model HIDs 961 and 4423), 60-step Galaxy workflow compiled via Workflow Foundry, workflow invocation, and three custom UDTs (`hokken_go_enrichment`, `hokken_venn_comparison`, and `hokken_revigo_summary`). |
| **Ruiz et al. (2021)**<br>*Anopheles gambiae* ATAC-seq & RNA-seq<br>(GSE152924) | bbd44e69cb8906b5e87b8ac73fb252a5<br>bbd44e69cb8906b57c33c7b295ddf477<br>bbd44e69cb8906b51645d526a9bd48d6<br>32f57ede003070c3,<br>6c3ed747eba652a6 | Primary ATAC-seq peak-calling, two-sided 51% overlap, and `either_frac` cross-fit history, raw ATAC QC history, companion RNA-seq quantification history, two published ATAC-seq workflows, and five custom UDTs (`ruiz_tn5_fragment_extractor`, `ruiz_macs2_212_callpeak`, `ruiz_reproducible_ths`, `ruiz_either_frac_shared_peaks`, and `ruiz_either_frac_crossfit`). |
| **Wang et al. (2024)**<br>*Aedes aegypti* scRNA-seq<br>(SRR26591148–49) | https://usegalaxy.org/u/scottcain/p/wang-2024-aaegypti-midgut-scrna-seq-reanalysis<br>https://usegalaxy.org/u/scottcain/h/wang-aaegypti-scrna-seq-veupathdb-annotation-rerun | Published Galaxy Page report and companion history containing STARsolo matrices, DropletUtils cell calls, custom MT_-prefixed GTF, Scanpy AnnData objects, and sample-blocked edgeR pseudobulk tables. |

**Table S4:** Computational artifacts from the spatial transcriptomics reproduction study. Galaxy histories, workflows, UDT definitions, and execution traces archived on UseGalaxy.org and GitHub.

| Artifact Type | Identifier / Link | Description |
| --- | --- | --- |
| Primary History | bbd44e69cb8906b596a6dfeaa66d7b35 | Reproduction of published *Acta2* 1,000-cell tutorial subset |
| Full Dataset History | bbd44e69cb8906b5e19946d9d8770575 | Full 33-gene spatial evaluation on Dataset 15 (public) |
| Single-Gene Workflow | fde4525ee8a0c99f | Executable workflow for target gene prediction |
| Full Evaluation Workflow | 63012fffa29eb7bb | Published 10-fold cross-validation workflow across 33 genes |
| Workflow Invocation | 7e806872876f7fb0 | Terminal execution trace for *Acta2* prediction |
| User-Defined Tools | `tissue_spatial_prediction v1.2.0`, `tissue_full_evaluation v1.0.0` | Containerized TISSUE UDT specifications |
| Interactive Pages | 14ebc62e2d1afb49, 787090c6bb87743a | Research notebook and evaluation report on UseGalaxy.org |
| Code Repository | https://github.com/nekrut/spatTr0rbit | Scripts, local references, and workflow templates |

**Table S5:**
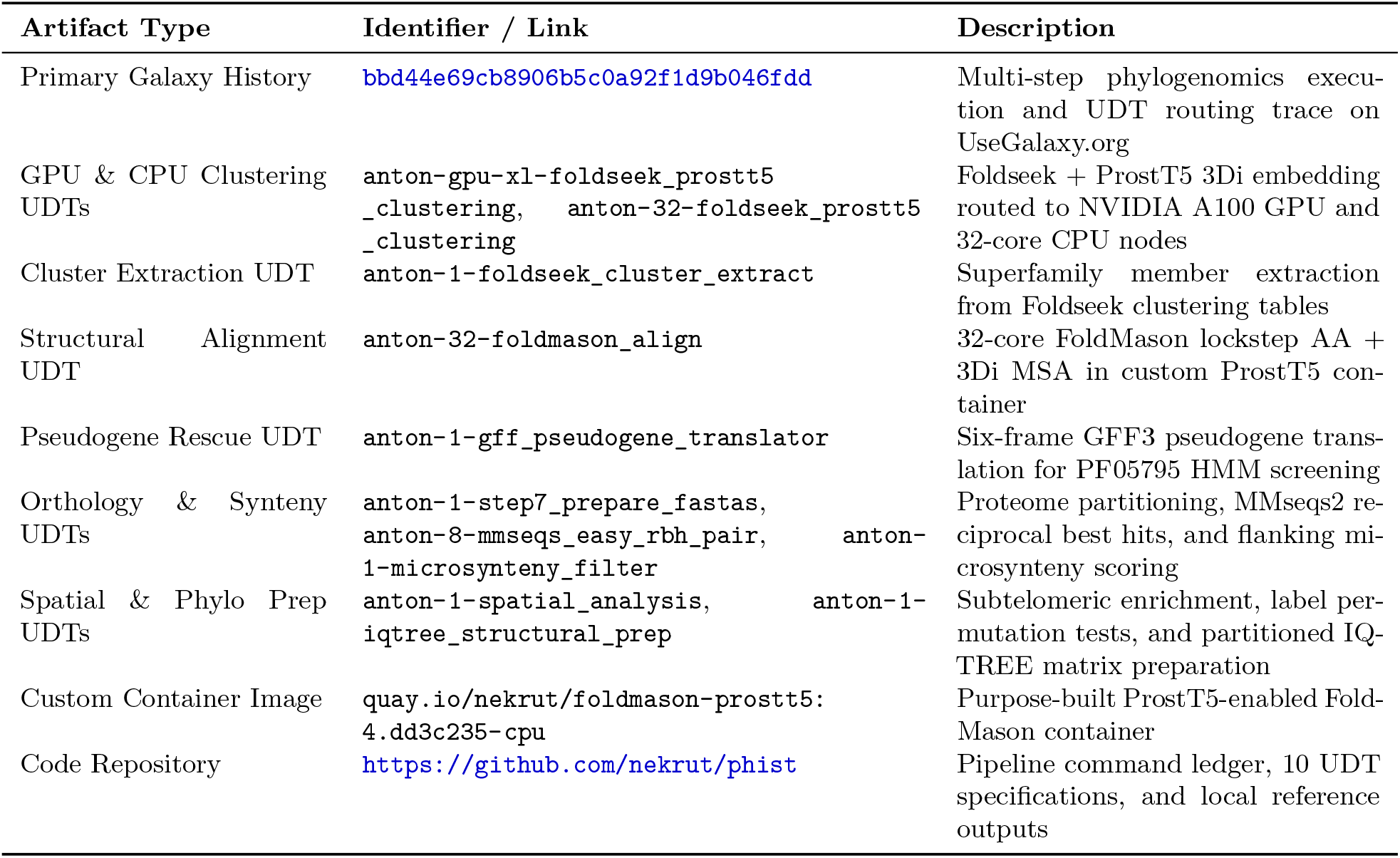
Computational artifacts from the *Plasmodium vivax* phylogenomics study. Galaxy histories, ten User-Defined Tool definitions, custom container images, and verification suites archived on UseGalaxy.org, Quay.io, and GitHub.

**Table S6:** Computational artifacts from the 42-pathogen molecular clock benchmark study. Galaxy histories, User-Defined Tool definitions, dataset collections, and published interactive reports archived on UseGalaxy.org and GitHub.

| Artifact Type | Identifier / Link | Description |
| --- | --- | --- |
| Preparation History | bbd44e69cb8906b5c7c1fd5515b95301 | Input validation and collection preparation for 42 cohorts |
| Analysis History | bbd44e69cb8906b55342b6485eac77f7 | End-to-end tree-free dating and diagnostics execution |
| Originals Archive | f9cad7b01a472135e65a92a49dc98cbe | Public binary archive of 42 author BEAST XML files |
| Batch Preparator UDT | `chronaeon_batch_preparator v1.0.0` | XML parser, FASTA extractor, and date validator |
| Cohort Analysis UDT | `chronaeon_cohort_analysis v3.2.2` | Tree-free dating, spline clock fitting, and LOOCV validation |
| Input Collections | e2f9a561d7c485b5 (XML), 25b287f1e09c05f2 (FASTA), 73f4b0a869dae7a4 (dates) | Synchronized 42-element BEAST XML, aligned FASTA, and tip-date collections |
| Output Collections | eacb6fa1da5e5cbe, c2c3f23ed16816c8 | Dating JSON and diagnostic plot collections for 42 cohorts |
| Interactive Page | feee8a2c33d4f9e1 | Published compendium evaluation report and concordance dashboard |
| Code, Prompts & Provenance | https://github.com/veg/chronaeon, `manuscript/prompts/`, `manuscript/figures/data/chronaeon/` | Verbatim Orbit harvest-ing/analysis prompts (`prompt.md`, `prompt-galaxy.md`, `prompt-galaxy-analysis.md`), benchmark TSVs, and comparator provenance table (`chronaeon_42_cohorts_provenance.tsv`) |

**Table S7:**
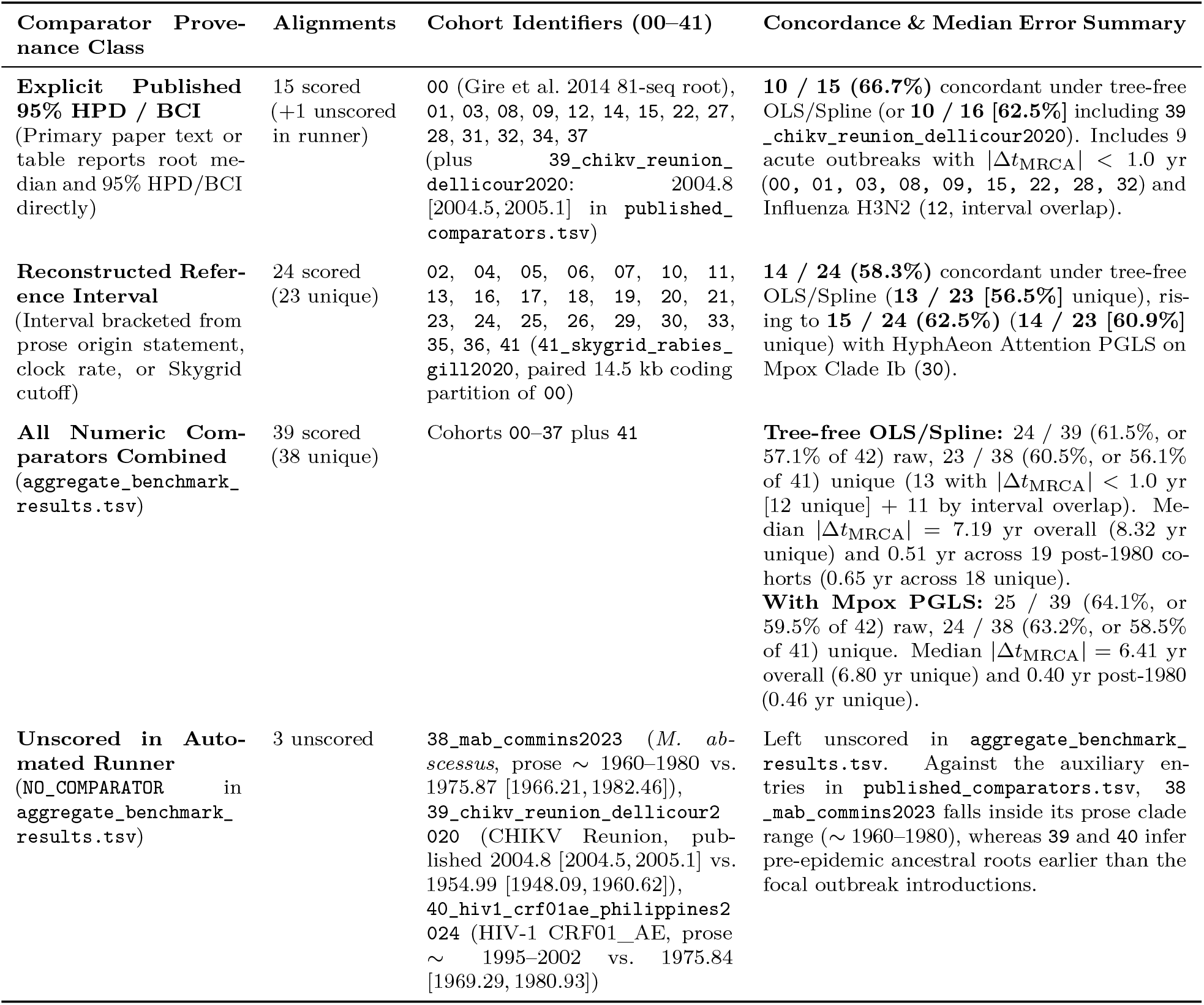
Provenance classification and concordance breakdown of reference comparators across the 42 ChronAeon pathogen benchmark alignments. Per-cohort estimates, Fieller 95% confidence intervals, comparator sources, and single-pass versus multi-model values are archived in `manuscript/figures/data/chronaeon/chronaeon_42cohorts_provenance.tsv`.

## Footnotes

1 Because executing arbitrary user-supplied tool definitions on public supercomputing infrastructure carries security and resource risks, User-Defined Tool registration currently requires account approval on UseGalaxy.org or routes through an automated packaging and review pipeline on UseGalaxy.eu (see Discussion).

